# Long-term field experiment on the impacts of the neonicotinoid dinotefuran and the organophosphate fenitrothion on a honeybee colony

**DOI:** 10.1101/014795

**Authors:** Toshiro Yamada, Yasuhiro Yamada, Kazuko Yamada, Hiroko Nakamura

**Author notes:** Authors’ addresses: Toshiro YAMADA (corresponding author,), Division of Material Science, Graduate School of Natural Science & Technology, Kanazawa University, Kakuma-machi, Kanazawa 920-1192, Japan; present address as an professor emeritus of Kanazawaa University: 2-10-15 Teraji, Kanazawa, Ishikawa 921-8178, Japan. Yasuhiro YAMADA, Department of Applied Physics, Graduate School of Engineering, University of Tokyo, 7-3-1 Hongo, Bunkyo-ku, Tokyo 113-8656, Japan. Kazuko YAMADA, present address: 2-10-15 Teraji, Kanazawa 921-8178, Japan. Hiroko Nakamura, Division of Material Science, Graduate School of Natural Science & Technology (Present: General Affairs Department), Kanazawa University, Kakuma-machi, Kanazawa 920-1192, Japan.

## Abstract

**Summary:** Neonicotinoides are persistent and highly toxic pesticides that have become popular instead of organophosphates, being suspected to be a trigger of massive disappearance of bees that raises concern in the world. The evaluation of the long-term influence for a whole colony in the natural environment is, however, not established yet. In this paper, we conducted a long-term field experiment and found different impacts on honeybee colonies *(Apis mellifera)* in an apiary between the neonicotinoid dinotefuran and the organophosphate fenitrothion even though whose concentrations in sugar syrup provided for bees were adjusted to have nearly equal short-term effects on a honeybee based on the median lethal dose (LD_50_) as well as the insecticidal activity to exterminate stinkbugs.

The colony with administration of dinotefran (dinotefuran colony) became extinct in 26 days, while the colony with administration of fenitrothion (fenitrothion colony) survived the administration for the same period. Furthermore, the fenitrothion colony succeeded to be alive for more than 293 days after administration, and also succeeded an overwintering, which indicates that colonies exposed to fenitrothion can recover after the exposure.

Meanwhile, the dinotefuran colony became extinct even though the intake of dinotefuran was estimated to be comparable with that of fenitrothion in terms of the LD_50_ of a honeybee. Moreover, the colonies in our previous long-term experiments where dinotefuran with higher concentration were administered only for first few days (Yamada et al., 2012) became extinct in 104 days and 162 days, respectively. From these results, we speculate that colonies exposed to dinotefuran hardly recover from the damage because dinotefuran is much more persistent than fenitrothion and toxic foods stored in cells can affect a colony in a long period.

## Introduction

Massive losses of honeybee colonies is becoming a worldwide problem (Van Engelsdorp et al., 2011; van Engelsdorp et al., 2012; Spleen et al. 2013; Steinhauer et al. 2014; van der Zee et al., 2012; van der Zee et al., 2014; Pirk et al., 2014). Many researchers have tried to find out the cause of them and have proposed various causes such as pesticides, mites, pathogens and so on. Recently pesticides, especially neonicotinoid pesticides which are persistent, systemic and high toxic, are strongly suspected of causing the massive losses based on many laboratory experiments and several longterm field experiments (van der Sluijs, 2013). Neonicotinoid pesticides (neonicotinoids) have been widely used in the world at present, even after a moratorium in the EU on the use of three neonicotinoids (imidacloprid, clothianidin, thiamethoxan) under the given limitations. In 2013, many papers have been reported on the adverse effects of neonicotinoids on insects (Prisco, 2013; EFSA, 2013a,b,c,d; Hatjina et al., 2013; Hunt & Krupke, 2013), mammals (EFSA, 2013e; Bal et al., 2013) and human (Taira et al., 2013).

A neonicotinoid has been evaluated by the LD_50_ (50% lethal dose) which is one way to measure the short-term poisoning potential (acute toxicity) of a material. This value can give the useful information on the acute toxicity in a short-term dose but cannot evaluate the chronic toxicity in a long-term one. In order to elucidate the anomalous behaviors of honeybee colony such as a colony collapse disorder and a failure in wintering, the impact of chronic toxicity on a honeybee colony in the fields is more important than that of acute toxicity realistically.

Field experiments include many uncontrollable factors such as honeybee behavior, weather, hornet attacks, mites and pathogens and so on. However, supposing that field experiments are conducted under the same circumstances, it becomes important to evaluate the honeybee behavior of an experimental colony because the other factors are generally offset by a control colony. As the behaviors of honeybees are uncontrollable and closely related to each other as eusocial insects in the fields, the experimental results in controllable laboratory testing under certain limited and special circumstances cannot be always applied to those in filed testing. In addition to this, when comparing the LD_50_ with the pesticide amount taken by an experimental colony in field testing, it should be considered that honeybees prefer pesticide-free nectar and natural pollen to sugar syrup and artificial pollen containing a pesticide.

According to our previous works (Yamada *et al.*, 2012; Yamada *et al.*, under submission), we have confirmed that high concentrations in sugar syrup (dinotefuran of 10 ppm; clothianidin of 4 ppm), which is used also to make pollen paste after mixing it with pollen, collapsed the honeybee colonies due to acute toxicity, low pesticide concentrations in sugar syrup (dinotefuran of 1 ppm; clothianidin of 0.4 ppm) collapsed the colonies due to chronic toxicity after having assumed the appearance of a colony collapse disorder (CCD) or an failure in wintering, and middle concentrations in sugar syrup (dinotefuran of 2 ppm; clothianidin of 0.8 ppm) damaged the colonies due to acute toxicity at the start of administration and due to chronic toxicity at the later period after having assumed the appearance of CCD and finally collapsed them.

It was confirmed that honeybees took toxic foods (sugar syrup, pollen paste) in the hive even when they could freely take nontoxic nectar from fields. Even though the low pesticide concentrations in our previous studies would cause the instantaneous death of honeybees due to acute toxicity judging from the LD_50_, in actual fact, the low concentrations hardly caused any instantaneous death. This result seems to be ascribed to the dilution of toxic sugar syrup or pollen paste in a beehive by pesticide-free nectar or natural pollen existing in the fields. The dilution ratio of toxic sugar syrup or pollen paste by pesticide-free nectar or natural pollen selectively taken from fields depends on the weather. These suggest that it is quite inadequate to assume that the field experimental conditions can be determined from the results of laboratory testing.

Recently,Pilling *et al.* (2013) reported that no detrimental effects on colony survival and overwintering success could be found at four-year repeated field exposures of thiamethoxam to pollen and nectar. However, the experimental concentrations of thiamethoxam are much lower than the residue concentration (53 ppb in pollen) reported by Johnson *et al.* (2010) and can be probably too low to affect a colony even due to its chronic toxicity. Incidentally, the actual average year-round concentration of a pesticide included in stored honey on a comb is unclear and the cumulative total intake of pesticide per bee is unknown in the report by Pilling *et al.*, 2013. Further, the result by Pilling **et al.** (2013) may be attributable to the dilution of poisoning pollen and nectar fed to a colony with nontoxic ones from fields, or only a very slight intake of toxic honey or pollen fed to a colony by honeybees.

Yamada *et al.* (2012) have conducted the field experiment at low, middle and high concentrations of neonicotinoids (dinotefuran, clothianidin) and Yamada **et al.** (under submission) have done at low and high concentrations of dinotefuran. So far, low and high concentration field-experiments of dinotefuran have been conducted twice by the authors (Yamada *et al.*, 2012, Yamada *et al.*, under submission) whose results have been replicated respectively but middle one has been done only once (Yamada *et al.*, 2012). Our previous results have revealed through a long-term field experiment that neonicotinoids lead to the gradual extinction of a honeybee colony due to chronic toxicity after the occurrence of many instantaneous honeybee-deaths at high concentration, some ones at middle concentration and no ones at low concentration due to acute toxicity in the beginning of experiment. The colony exposed to neonicotinoids dwindled away to nothing after showing an aspect of CCD or failed in overwintering.

Dinotefuran and fenitrothion are known as a representative pesticide of neonicotinoids and that of organophosphates in Japan. Though the impact of fenitrothion on birds, insects, fish, honeybees and so on and the persistent residues in the environment has been widely investigated by long-term field monitoring (Mitchelland Roberts, 1984), it is uncertain whether fenitorothion causes CCD or not. In this work we will confirm whether these results obtained from neonicotinoids can be applied to organophosphates such as fenitrothion or not. Here, we will elucidate the impact of fenitrothion on a honeybee colony during long-term exposure to a pesticide comparing it with dinotefuran. In this work we will clarify the followings. (1) Will which pesticide of the neonicotinoid dinotefuran and the organophosphate fenitrothion become extinct faster after both pesticides are prepared to be the identical insecticidal activity for stinkbugs considering actual usage in Japan? Which has actually higher toxicity for a honeybee colony? (2) How will each colony behave when we feed toxic sugar syrup which is newly prepared every observation? What difference in behavior of a honeybee colony can be caused between dinotefuran and fenitrothion? (3) How will the surviving colony behave after it is damaged by the pesticide when it continues to take nontoxic sugar syrup instead of toxic sugar syrup just after either colony has become extinct? How much will the stored toxic sugar syrup (honey) in the hive continue to affect the honeybee colony?

## Materials and Methods

### Ethics statement

We clearly state that no specific permissions were required for these locations/activities because the apiary at which we performed the experiments for this study belongs to the author (Toshiro Yamada). We confirm that the field studies did not involve endangered or protected species.

### Materials and preparation of pesticide concentrations

Experiments were performed in 2012 to 2013 under experimental conditions as tabulated in Table 1. STARCKLE MATE^®^ (10% dinotefuran; Mitsui Chemicals Aglo, Inc., Tokyo, Japan) and SUMITHION EMULSION (50% fenitrothion; Sumitomo Co. Ltd., Osaka, Japan) used in this study. On comparing the effect of both pesticides on honeybees, we adopted the concentrations to exterminate stinkbugs considering general usage of Japan and a very wide range of the LD_50_ which each pesticide has. Each concentration of dinotefuran and fenitrothion was determined at the one-fiftieth of the spraying concentration (100ppm for dinotefuran, 500 ppm for fenitrothion) to exterminate stinkbugs by referring our previous results, which was dinotefuran of 2 ppm and fenitrothion of 10 ppm, respectively. Neonicotinoids of dinotefuran and clothianidin, which are adjusted to have a same insecticidal activity affecting stinkbug, are confirmed to have almost the same effect on honeybees. The concentrations of dinotefuran and clothianidin caused some instant honeybee-deaths at the beginning and afterwards the gradual extinction of a honeybee colony after giving the appearance of CCD when they are administered into colonies through both sugar syrup and pollen paste (Yamada *et al.*, 2012).

**Table 1.**
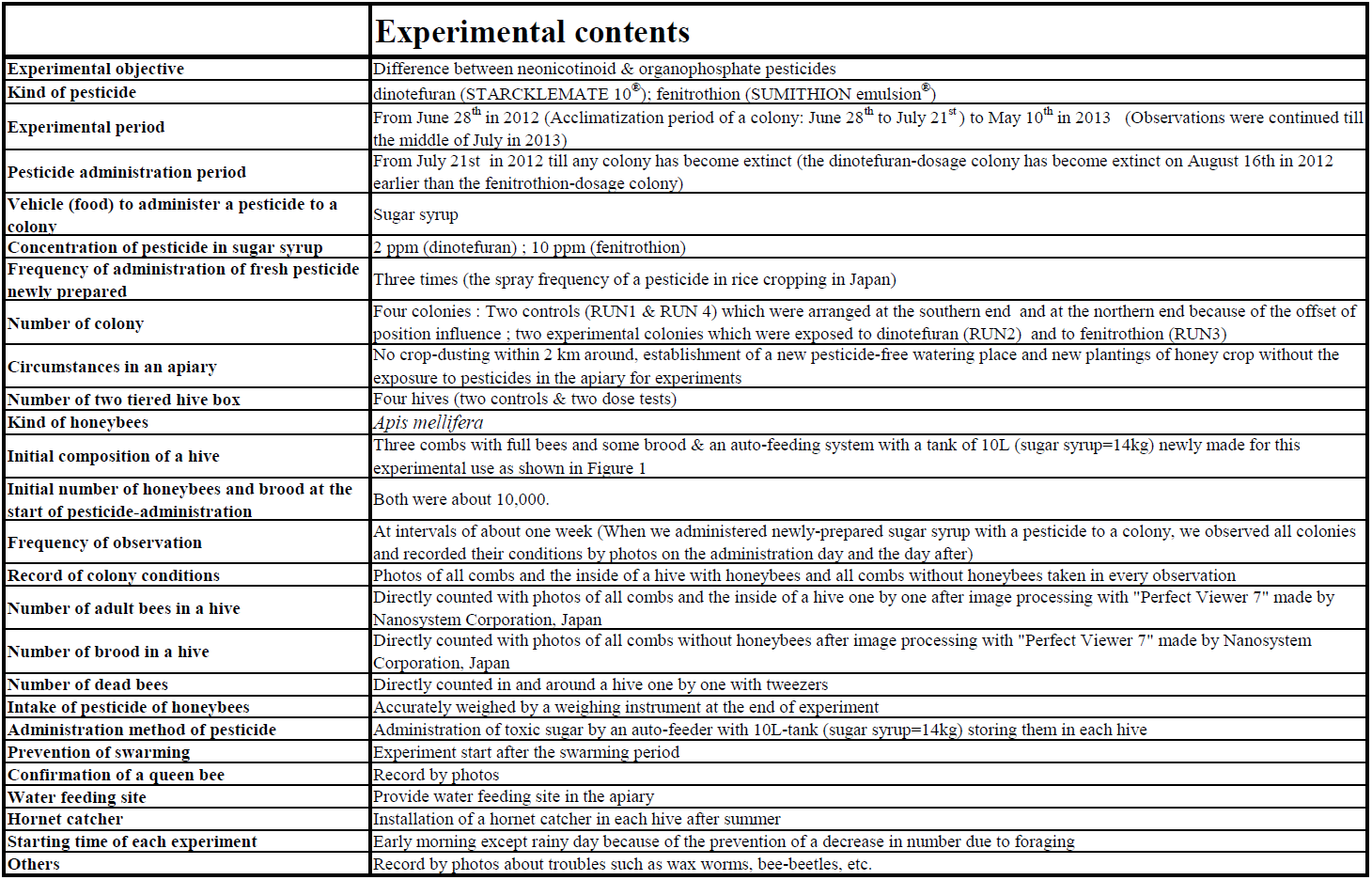
Outline of experimental conditions in this work

|  | Experimental contents |
| --- | --- |
| Experimental objective | Difference between neonicotinoid & organophosphate pesticides |
| Kind of pesticide | dinotefuran (STARCKLEMATE 10<sup>®</sup>); fenitrothion (SUMITHION emulsion<sup>®</sup>) |
| Experimental period | From June 28<sup>th</sup> in 2012 (Acclimatization period of a colony: June 28<sup>th</sup> to July 21<sup>st</sup>) to May 10<sup>th</sup> in 2013 (Observations were continued till the middle of July in 2013) |
| Pesticide administration period | From July 21<sup>st</sup> in 2012 till any colony has become extinct (the dinotefuran-dosage colony has become extinct on August 16<sup>th</sup> in 2012 earlier than the fenitrothion-dosage colony) |
| Vehicle (food) to administer a pesticide to a colony | Sugar syrup |
| Concentration of pesticide in sugar syrup | 2 ppm (dinotefuran) ; 10 ppm (fenitrothion) |
| Frequency of administration of fresh pesticide newly prepared | Three times (the spray frequency of a pesticide in rice cropping in Japan) |
| Number of colony | Four colonies : Two controls (RUN1 & RUN 4) which were arranged at the southern end and at the northern end because of the offset of position influence ; two experimental colonies which were exposed to dinotefuran (RUN2) and to fenitrothion (RUN3) |
| Circumstances in an apiary | No crop-dusting within 2 km around, establishment of a new pesticide-free watering place and new plantings of honey crop without the exposure to pesticides in the apiary for experiments |
| Number of two tiered hive box | Four hives (two controls & two dose tests) |
| Kind of honeybees | *Apis mellifera* |
| Initial composition of a hive | Three combs with full bees and some brood & an auto-feeding system with a tank of 10L (sugar syrup=14kg) newly made for this experimental use as shown in Figure 1 |
| Initial number of honeybees and brood at the start of pesticide-administration | Both were about 10,000. |
| Frequency of observation | At intervals of about one week (When we administered newly-prepared sugar syrup with a pesticide to a colony, we observed all colonies and recorded their conditions by photos on the administration day and the day after) |
| Record of colony conditions | Photos of all combs and the inside of a hive with honeybees and all combs without honeybees taken in every observation |
| Number of adult bees in a hive | Directly counted with photos of all combs and the inside of a hive one by one after image processing with "Perfect Viewer 7" made by Nanosystem Corporation, Japan |
| Number of brood in a hive | Directly counted with photos of all combs without honeybees after image processing with "Perfect Viewer 7" made by Nanosystem Corporation, Japan |
| Number of dead bees | Directly counted in and around a hive one by one with tweezers |
| Intake of pesticide of honeybees | Accurately weighed by a weighing instrument at the end of experiment |
| Administration method of pesticide | Administration of toxic sugar by an auto-feeder with 10L-tank (sugar syrup=14kg) storing them in each hive |
| Prevention of swarming | Experiment start after the swarming period |
| Confirmation of a queen bee | Record by photos |
| Water feeding site | Provide water feeding site in the apiary |
| Hornet catcher | Installation of a hornet catcher in each hive after summer |
| Starting time of each experiment | Early morning except rainy day because of the prevention of a decrease in number due to foraging |
| Others | Record by photos about troubles such as wax worms, bee-beetles, etc. |

Incidentally, focusing on the LD_50_, the LD_50_ values of dinotefuran and clothianidin widely ranges from 7.6 ng/bee (US-EPA, 2004) to 75 ng/bee (Iwasa *et al.*, 2004) and from 20 ng/bee (US-EPA, 1995) to 380 ng/bee (US-EPA, 1995), respectively. The average of a minimum and a maximum of each LD_50_ is about 41 ng/bee for dinotefuran and 200 ng/bee for fenitrothion. Judging from the ratio of these averages which is about five, 2 ppm of dinotefuran and 10 ppm of fenitrothion, which have the same insecticidal activity to exterminate stinkbugs, can be estimated to have almost the same insecticidal activity in terms of the LD_50_ for a honeybee.

As the frequency of spraying of a pesticide (dinotefuran, fenitrothion) is usually about three times in order to exterminate stinkbugs in rice cropping in Japan, we have determined to administer fresh pesticides newly prepared three times. We observed the colonies and got a photographic record of them (all combs with and without honeybees and the inside of a beehive and the outside just before the administration of a fresh pesticide and the day after in order to investigate an acute toxic effect of insecticidal activity of a pesticide. Comparing the numbers of adult bees and dead bees the day after the new administration of the pesticide with those about one week after, we examined the toxicity change of the administered pesticide.

The experimental concentrations of these pesticides were realistic in the field of Japan from the facts that the concentration of clothianidin near rice paddies was about 5 ppm (Kakuta *et al.*, 2011) and maximum residue limits (MRLs) of agricultural chemicals in foods in Japan (JFCRF, 2014). Then the experimental concentration of dinotefuran was determined from the insecticidal activity of clothianidin to the honeybee was about 2.5 times as much as that of dinotefuran; namely, dinotefuran of 10 ppm is equivalent to clothianidin of 4 ppm (Yamada *et al.*, 2012).The insecticidal activity of dinotefuran for honeybees was almost equivalent to that of chrothianidin after equalizing their insecticidal activity for stinkbugs. And the pesticides administered to the colonies in this field experiment seem to be diluted with pesticide-free nectar collected from the fields by foraging bees. According to the information from beekeepers, bees generally prefer to consume nectar and their own honey, so the consumption of feed (sugar syrup) indicates a lack of these and then while being fed, bees will consume some feed and store some (ColonyMonitoring.com, 2012). Incidentally, orange blossom honey contained acetamiprid of 0.05 ppm in Japan (Notice from Tamagawa Gakuen, 2013) while acetamiprid is usually sprayed on oranges in the concentration of about 60 ppm in Japan.

### Methods used in field experiments

Four beehives, each with 3 numbered combs and a feeder, were sited facing east on a hill. They were aligned in order of RUN number; the control colony (RUN1), the dinotefuran-dosage one (RUN2), the fenitrothion-dosage one (RUN3) and the control one (RUN4) from the south to the north. Two controls were arranged at both ends because of the confirmation of difference between north and south.

Pesticide-free sugar syrup was fed into every colony from June 28^th^ in 2012 to the early morning of July 21^st^ as a preliminary experiment in order to acclimatize the colonies to the experimental apiary after the swarming season. After the period of acclimatization, we administered each pesticide into the dinotefuran colony (RUN2) and the fenitrothion colony (RUN3), respectively, till either colony became extinct while each toxic sugar syrup with a pesticide was replaced with newly-prepared (fresh) one every administration date. After an experimental colony became extinct, we exchanged toxic sugar syrup with pesticide-free one in the surviving experimental colony in order to investigate whether the surviving colony exposed to the pesticide can recover from the damage of a pesticide or not.

We observed all colonies and took photos of all combs with bees, those without bees, the inside with residual bees of each hive box, surrounding circumstances and so on about every week on the administration day and the day after. The total number of adult bees on all combs, which were numbered and ordered numerically in every hive, and a feeder and the inside of the hive box (4 walls and bottom) was counted directly and accurately from photographs (sometimes enlarged) of all combs with “Perfect Viewer 7” made by Nanosystem Corporation, Japan. The total number of capped brood was counted in a similar manner, after directly shaking the bees off each comb as shown in Figure 1. Though we have tried to develop a new automatic counting software with binarizing photo images, we cannot succeed in accurate counting of them because it cannot accurately count overlaid bees, bees and capped brood on blurred image, those on low contrast one or those on low brightness one even when changing the threshold. To obtain the total number of dead bees in and around the hive and feeder, the hive was placed on a large tray. The total number of dead bees in the tray, feeder, and hive was counted directly, one by one with a pair of tweezers.

**Figure 1.**
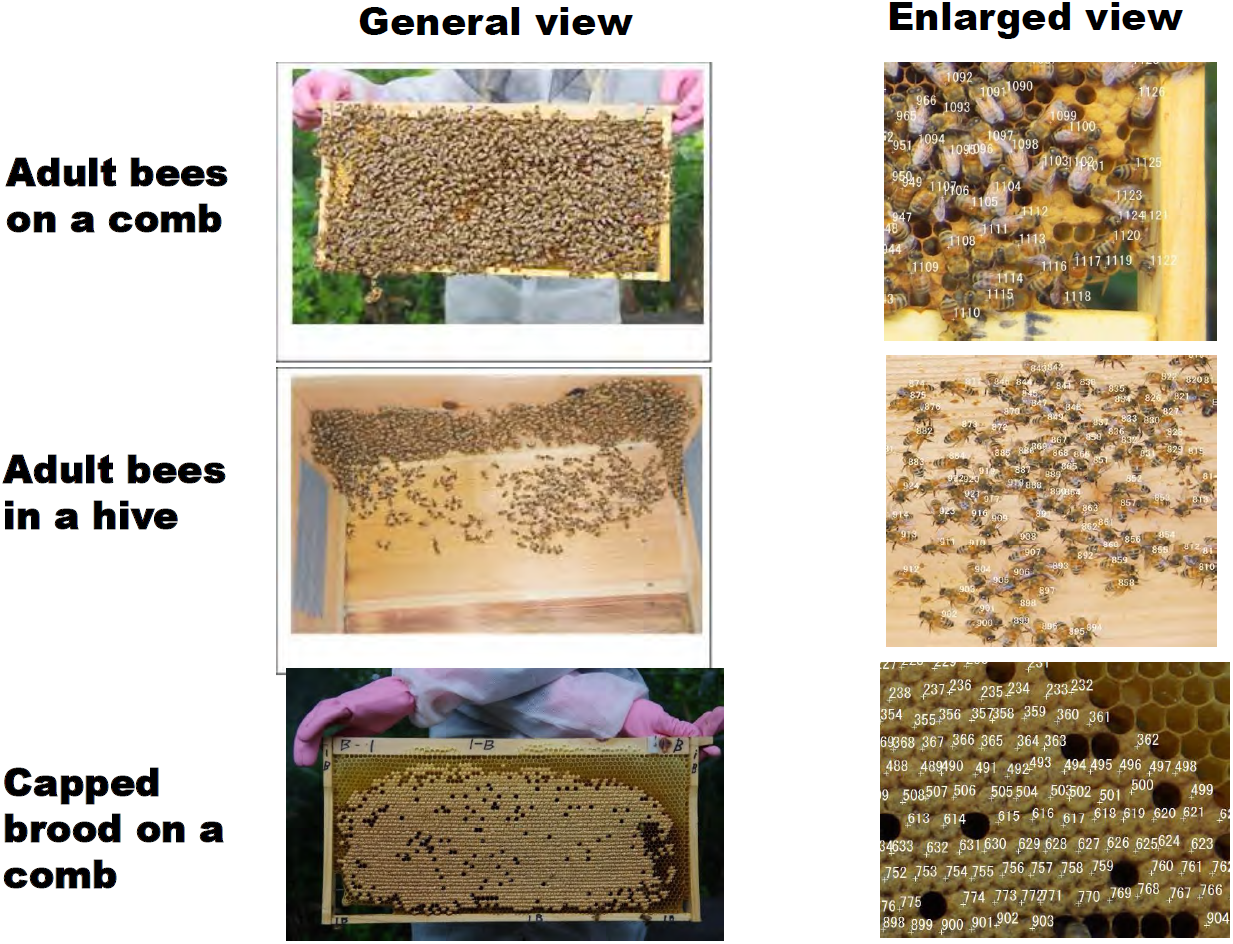
Counting method of adult bees and capped brood in a hive. We counted almost all of adult bees and capped brood in the hive with numbering bees and capped brood on a photo by a numbering software, Nanosystem Corporation, Japan, which we have taken on the early morning before foraging bees went out.

The queen bee in the hive was photographically recorded on each measurement date, as were specific situations such as the presence of chalk brood or wax moth larvae and the evidence of Asian giant hornet attacks. During the experimental period, hive status was recorded at intervals of 1 h with a digital camera.

We performed the experiment early in the morning on fine or cloudy days, before the foraging bees left the hive from June 28^th^ in 2012 to May 10^th^ in 2013. We continued to observe the pesticide-free colonies till the middle of July in 2013 after finishing this experiment (May 10^th^ in 2013) in order to clarify the normal behavioral standards of a honeybee colony for a year.

In order to decrease in unclearness and diversity of uncontrollable factors contained in field experiments, we selected an experimental site where there are not any aerial-sprayed paddy fields and orchards in the vicinity. We located a honeybee-watering place in the experimental apiary to supply pesticide-free water and planted leaf mustard *(Brassica juncea)* and hairy vetch *(Vicia villosa)* in the experimental site to prevent honeybees from taking nectar and pollen contaminated by pesticides in order to minimize the effects of environmental factors.

The consumption of sugar syrup by honeybees was accurately measured by a weighing instrument having an accuracy of 0.1g in every observation. The net intake of a pesticide was obtained from the amount of sugar syrup consumed by honeybees. The cumulative total intake of each active ingredient (dinotefuran, fenitrothion) was obtained from the amount of sugar syrup consumed by honeybee colony during the pesticide-administration period. The interval intake of a pesticide by a colony between two observation dates (a certain observation date and the previous one) was obtained from the consumption of sugar syrup with a pesticide. The intake of a pesticide per bee was estimated from dividing the cumulative total intake of the pesticide in a colony by the sum of the total number of newborn honeybees, the number of initial honeybees and that of the capped brood at the colony extinction.

Strictly speaking, this experiment cannot be always conducted under the very same conditions as the natural environment near an actual apiary, because sugar syrup is not same as nectar in fields and the feeding area in this work is not the same as that in an actual apiary. That is, honeybees in an experimental colony of this work take not only toxic sugar syrup in a hive but also nectar which is controlled mainly so as to be nontoxic by pesticide-free flowers in an apiary, while those in a colony of an actual apiary take nectar which is toxic and/or nontoxic in fields. In addition, not only foraging bees but also house bees may take sugar syrup in this work, while only foraging bees take nectar in fields in an actual apiary. Despite these differences from an actual apiary, we believe that this experiment can possibly replicate most of the phenomena occurring in an actual apiary though we have to pay attention to them.

## Results

### Long-term observations

The experiment was conducted under the nearly natural environment where honeybees can freely take foods in fields if they do not like to take toxic sugar syrup in a hive. We found that the dinotefuran colony (RUN2) became extinct but the fenitrothion colony (RUN3) survived on August 16^th^ and thereafter in the subsequent recovery experiment the fenitrothion colony continued to survive after it succeeded in overwintering. Details of observations are as follows:

In the acclimatization period from June 28^th^ to July 21^st^ in 2012, the somewhat different numbers of adult bees and capped brood among colonies on June 28^th^ became almost the same on July 21^st^ when the pesticide-administration experiment started after we had taken photographs of all of combs with and without honeybees and the honey bees left behind in every hive box with combs being removed.

We started to administer each pesticide (dinotefuran, fenitrothion) into the colony on July 21^st^ and continued to do till August 16^th^ when the dinotefuran colony (RUN2) became extinct but the fenitrothion colony (RUN3) survived. In the administration period of pesticide, fresh sugar syrup with each pesticide newly prepared was fed into each colony three times, on July 21^st^, July 27^th^ and August 3^rd^. We discontinued the administration of fenitrothion and began to feed pesticide-free sugar syrup into the fenitrothion colony (RUN3) on August 16^th^ similarly to the control colonies (RUN1 and RUN4). The colony in which dinotefuran was administered (RUN2) rapidly dwindled away to nothing within a month from the start of pesticide administration, but the colony where fenitrothion was administered (RUN3) and both control ones (RUN1 & RUN4)) succeeded in overwintering without extinction. We judged that both control colonies and fenitrothion one succeded in overwintering on February 1^st^ in 2013. We administered a preventive medicine for foul brood following the instructions of Japan Beekeeping Association on March 17 in 2013. We finished the experiment on May 10^th^ in 2013 after good results of the foul brood test by the Livestock Health Center in Ishikawa Prefecture in Japan because the colonies became very vigorous. After that we continued to observe these three colonies (RUN1, 3 and 4) till the middle of July in 2013 for the investigation of the year round behavior of honeybee colony. The queen existed in every colony till the colony became extinct.

All the dinotefuran colonies where the neonicotinoid dinotfuran was administered have ended in extinction during the three long-term field experiments conducted from July of 2010 to May of 2013 with different courses depending on their concentration and administration period. On the other hand the fenitrothion colony dwindled during the administration of fenitrothion assuming a similar aspect to acute toxicity but it rapidly recuperated the vigor after the discontinuance of the administration. As a consequence, the fenitrothion colony succeeded in overwintering similarly to the control colony. It is desirable that this result is reproduced by other experiments as it was obtained from only one colony in this work.

### Measurement of number of dead bees

We measured the interval number of dead bees in an interval between two adjacent observation dates existing inside (on the bottom and in a feeder) and outside (mainly the front) of the beehive. Table 2 shows the interval number of dead bees at every observation date. These results were illustrated in Figure 2 after the conversion of the interval number of dead bees between two adjacent observations into the number of dead bees per day (daily number of dead bees). The followings can be seen from Table 2 and *Figure* 2:

**Figure 2.**
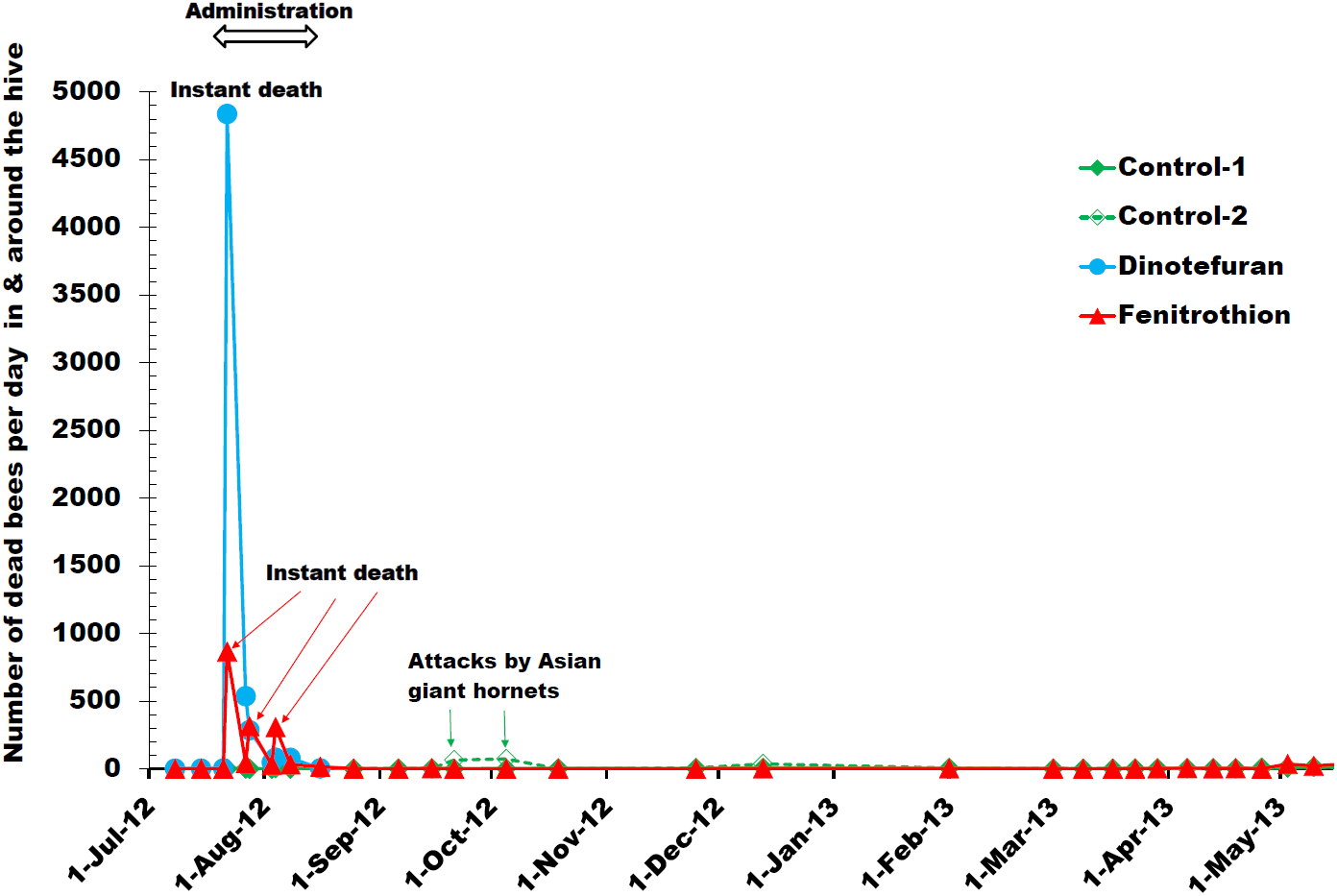
Daily number of dead bees. “Control 1, 2”, “DINOTEFURAN” and “FENITROTHION” indicate the colonies supplied with sugar syrup containing no pesticide, dinotefuran and fenitrothion, respectively. These pesticides were administered into their target colonies from July 21^st^ to August 16 in 2Q12. We defined a death of honeybees within a day after the administration of the pesticide (dinotefuran) as an instant death. The massive death of honeybees in Control-2 between September 21^st^ and October 5^th^ were supposed to be caused by the attacks of Asian giant hornets because we found dead Asian giant hornets and alive ones in front of the hive.

**Table 2.** Interval number of dead bees

| **Date** | **RUN1**<br>**Control 1**<br>**without pesticide** | **RUN2**<br>**Dinotefuran**<br>**2 ppm** | **RUN3**<br>**Fenitorothion**<br>**10 ppm** | **RUN4**<br>**Control 2**<br>**without pesticide** | **Note** |
| --- | --- | --- | --- | --- | --- |
| 8-Jul-12 | 2 | 10 | 2 | 1 |  |
| 15-Jul-12 | 18 | 1 | 0 | 0 |  |
| 21-Jul-12 | 2 | 5 | 3 | 3 | Beginning of pesticide administration |
| 22-Jul-12 | 0 | 4838 | 865 | 0 | Instant death |
| 27-Jul-12 | 10 | 2682 | 216 | 3 |  |
| 28-Jul-12 | 0 | 284 | 314 | 0 | Instant death |
| 3-Aug-12 | 4 | 276 | 166 | 7 |  |
| 4-Aug-12 | 9 | 81 | 307 | 2 | Instant death |
| 8-Aug-12 | 2 | 318 | 132 | 1 |  |
| 16-Aug-12 | 6 | 23 | 120 | 6 |  |
| 25-Aug-12 | 0 |  | 16 | 1 |  |
| 6-Sep-12 | 2 |  | 1 | 32 |  |
| 15-Sep-12 | 24 |  | 34 | 7 |  |
| 21-Sep-12 | 0 |  | 2 | 389 | Attacks by Asian giant hornets |
| 5-Oct-12 | 14 |  | 6 | 1017 | Attacks by Asian giant hornets |
| 19-Oct-12 | 51 |  | 5 | 9 |  |
| 25-Nov-12 | 56 |  | 23 | 215 |  |
| 13-Dec-12 | 122 |  | 42 | 648 | Attacks by Asian giant hornets |
| 1-Feb-13 | 185 |  | 115 | 317 | Natural death in winter |
| 1-Mar-13 | 34 |  | 21 | 13 |  |
| 9-Mar-13 | 3 |  | 0 | 2 |  |
| 17-Mar-13 | 10 |  | 3 | 5 |  |
| 23-Mar-13 | 6 |  | 11 | 15 |  |
| 29-Mar-13 | 16 |  | 16 | 20 |  |
| 6-Apr-13 | 29 |  | 37 | 32 |  |
| 13-Apr-13 | 33 |  | 19 | 20 |  |
| 19-Apr-13 | 11 |  | 24 | 22 |  |
| 26-Apr-13 | 32 |  | 8 | 66 |  |
| 3-May-13 | 57 |  | 240 | 99 | Mainly drones |
| 10-May-13 | 99 |  | 143 | 100 | Attacks by Asian giant hornets |

In experimental colonies (RUN2 with dinotefuran & RUN3 with fenitrothion) many dead bees occurred just after the first administration of pesticide from July 21^st^ to 22^nd^. In RUN2 with dinotefuran more than half (52.7 percent) of initial adult bees were instantly killed, and in RUN3 with fenitrothion about one tenth (9.7 percent) of adult bees died instantly. Much more dead bees tended to occur just after the administration of pesticides newly prepared for the periods from July 21^st^, 27^th^ and August 3^rd^ to 22^nd^, 28^th^ and August 4^th^, respectively than for the subsequent periods from July 22^nd^, 28^th^ and August 4^th^ to 27^th^, August 3^rd^ and 8^th^, respectively. Especially, such a tendency was strongly in evidence for fenitrothion (RUN3). In control colonies (RUN1, RUN4), any dead bees hardly occurred except in cases of the attack by Asian giant hornets and of the death in overwintering.

### Measurement of number of adult bees and capped brood

Table 3 shows the numbers of adult bees and capped brood in this work. In this table, figures written in red denote values in administration periods of pesticides and the others do in pesticide-free periods. Figures 3 and 4 show the changes in the numbers of adult bees and capped brood, respectively. We can find that dinotefuran can affect adult bees much more adversely than fenitrothion with the same insecticidal activity for stinkbugs while both of the pesticides can affect brood adversely to about the same degree. Details are below:

**Figure 3.**
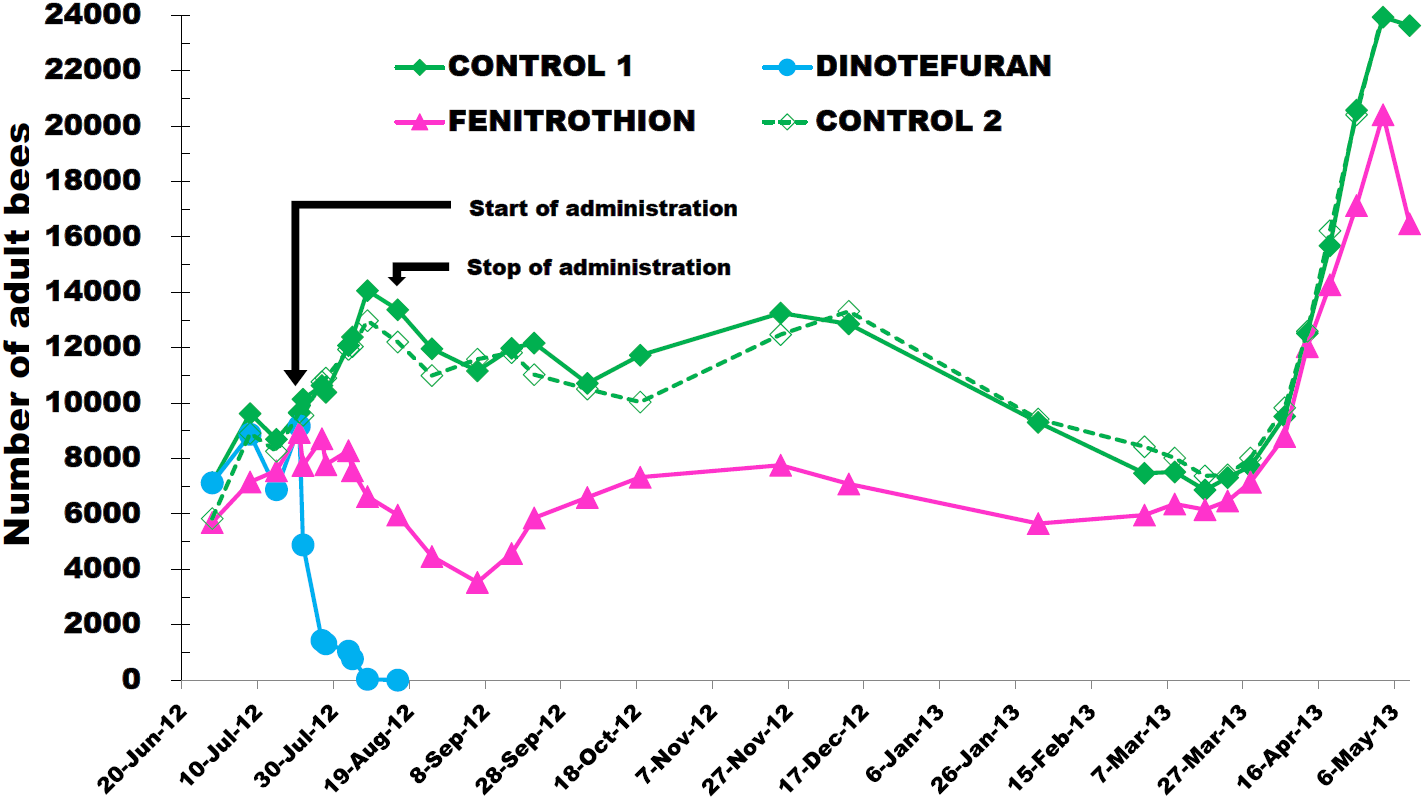
Change in the number of adult bees. “Control 1, 2”, “DINOTEFURAN” and “FENITROTHION” indicate the colonies supplied with sugar syrup containing no pesticide, dinotefuran and fenitrothion, respectively. These pesticides were administered into their target colonies from July 21^st^ to August 16 in 2012. The queen existed in every colony to the end of each experiment; that is, the queen in the dinotefuran colony existed till extinction.

**Figure 4.**
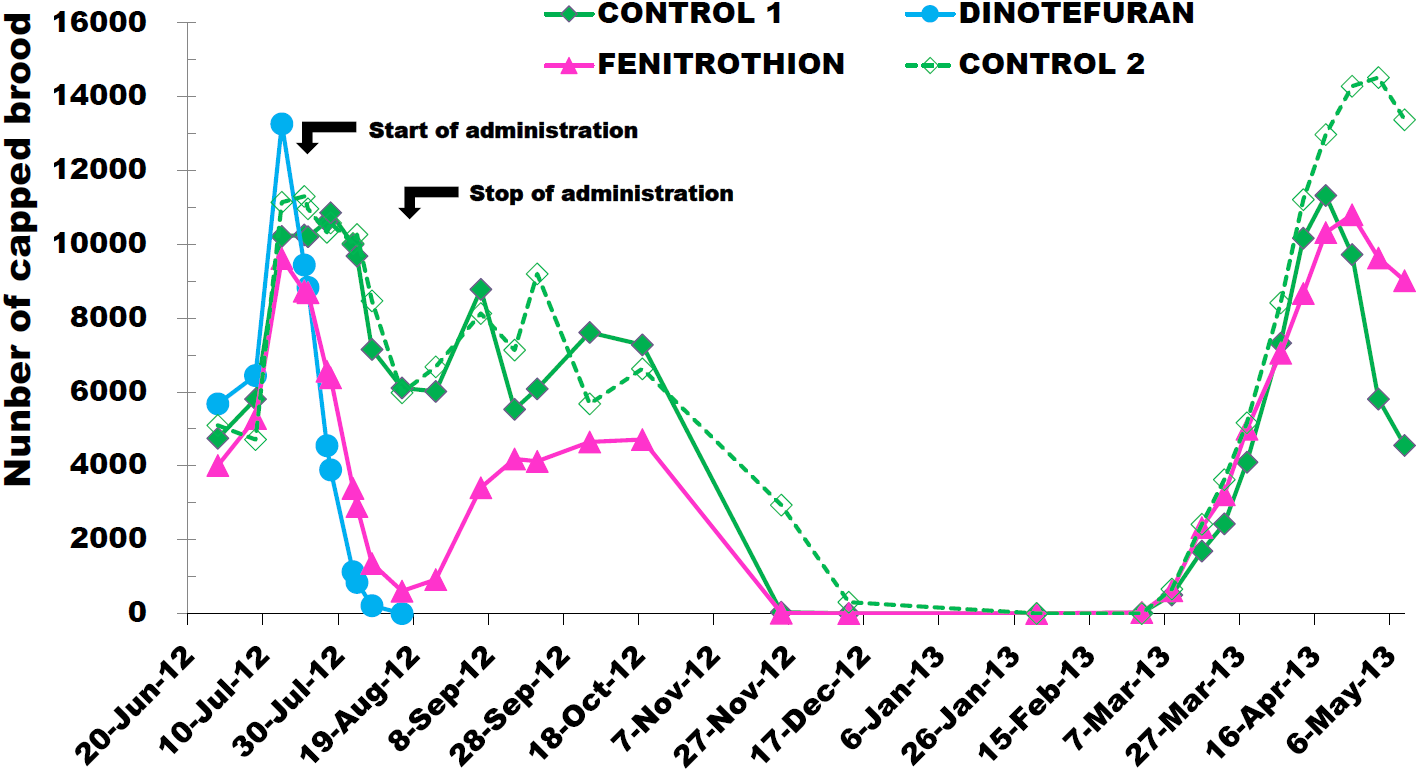
Change in the number of capped brood. “Control 1, 2”, “DINOTEFURAN” and “FENITROTHION” indicate the colonies supplied with sugar syrup containing no pesticide, dinotefuran and fenitrothion, respectively. These pesticides were administered into their target colonies from July 21^st^ to August 16 in 2012.

**Table 3.** Numbers of adultbees and capped brood

| Date | Elapsed days | Days from pesticide administration | RUN1 (Control 1) |  | RUN2 (Starcklemate) |  | RUN3 (Sumithion) |  | RUN4 (Control 2) |  |
| --- | --- | --- | --- | --- | --- | --- | --- | --- | --- | --- |
|  |  |  | without pesticide |  | dinotefuran : 2 ppm |  | fenitrothion : 10 ppm |  | without pesticide |  |
|  |  |  | Adult Bee | Capped Brood | Adult Bee | Capped Brood | Adult Bee | Capped Brood | Adult Bee | Capped Brood |
| 6/28/2012 | 0 | -23 | 7136 | 4746 | 7119 | 5679 | 5690 | 4012 | 5832 | 5094 |
| 7/8/2012 | 10 | -13 | 9621 | 5806 | 8877 | 6446 | 7157 | 5286 | 8917 | 4710 |
| 7/15/2012 | 17 | -6 | 8695 | 10215 | 6878 | 13267 | 7565 | 9619 | 8265 | 11143 |
| 7/21/2012 | 23 | 0 | 9647 | 10254 | 9173 | 9442 | 8943 | 8732 | 9665 | 11301 |
| 7/22/2012 | 24 | 1 | 10136 | 10210 | 4885 | 8834 | 7750 | 8694 | 9558 | 10967 |
| 7/27/2012 | 29 | 6 | 10633 | 10617 | 1434 | 4548 | 8721 | 6563 | 10770 | 10329 |
| 7/28/2012 | 30 | 7 | 10391 | 10858 | 1313 | 3891 | 7786 | 6389 | 10901 | 10581 |
| 8/3/2012 | 36 | 13 | 12083 | 10000 | 1049 | 1131 | 8289 | 3390 | 11939 | 10025 |
| 8/4/2012 | 37 | 14 | 12389 | 9687 | 771 | 840 | 7559 | 2901 | 12041 | 10269 |
| 8/8/2012 | 41 | 18 | 14065 | 7154 | 33 | 208 | 6625 | 1352 | 12978 | 8472 |
| 8/16/2012 | 49 | 26 | 13371 | 6111 | 0 | 0 | 5961 | 607 | 12207 | 5977 |
| 8/25/2012 | 58 | 35 | 11961 | 6014 |  |  | 4467 | 918 | 10997 | 6684 |
| 9/6/2012 | 70 | 47 | 11165 | 8783 |  |  | 3534 | 3406 | 11582 | 8126 |
| 9/15/2012 | 79 | 56 | 11980 | 5531 |  |  | 4576 | 4187 | 11825 | 7135 |
| 9/21/2012 | 85 | 62 | 12166 | 6086 |  |  | 5859 | 4119 | 11025 | 9202 |
| 10/5/2012 | 99 | 76 | 10715 | 7615 |  |  | 6593 | 4648 | 10510 | 5679 |
| 10/19/2012 | 113 | 90 | 11726 | 7280 |  |  | 7326 | 4713 | 10038 | 6628 |
| 11/25/2012 | 150 | 127 | 13255 | 36 |  |  | 7755 | 10 | 12477 | 2937 |
| 12/13/2012 | 168 | 145 | 12858 | 0 |  |  | 7080 | 0 | 13316 | 305 |
| 2/1/2013 | 218 | 195 | 9306 | 0 |  |  | 5652 | 0 | 9421 | 0 |
| 3/1/2013 | 246 | 223 | 7464 | 17 |  |  | 5957 | 26 | 8426 | 0 |
| 3/9/2013 | 254 | 231 | 7512 | 497 |  |  | 6358 | 619 | 8017 | 660 |
| 3/17/2013 | 262 | 239 | 6862 | 1691 |  |  | 6152 | 2329 | 7372 | 2419 |
| 3/23/2013 | 268 | 245 | 7312 | 2431 |  |  | 6470 | 3217 | 7416 | 3624 |
| 3/29/2013 | 274 | 251 | 7720 | 4097 |  |  | 7143 | 4994 | 8018 | 5165 |
| 4/7/2013 | 283 | 260 | 9518 | 7326 |  |  | 8797 | 7053 | 9833 | 8414 |
| 4/13/2013 | 289 | 266 | 12523 | 10166 |  |  | 12038 | 8670 | 12594 | 11211 |
| 4/19/2013 | 295 | 272 | 15677 | 11324 |  |  | 14275 | 10320 | 16221 | 12975 |
| 4/26/2013 | 302 | 279 | 20574 | 9725 |  |  | 17132 | 10803 | 20412 | 14287 |
| 5/3/2013 | 309 | 286 | 23935 | 5808 |  |  | 20413 | 9631 | 24100 | 14521 |
| 5/10/2013 | 316 | 293 | 23629 | 4551 |  |  | 16477 | 9020 | 27670 | 13380 |

(Note) Red numbers shows an administration period of pesticide.

The dinotefuran colony (RUN2) shows a drastic decrease of 46.7 percent in the number of adult bees within a day from July 21^st^ to July 22^nd^ in comparison with the initial number on July 21^st^. The decrease in the number of adult bees (4288; 46.7 percent) is somewhat less than the number of dead bees (4838; 52.7 %) in the same interval. This suggests that almost all of dead bees died on the spot considering the number of newborn adult bees within a day. The dinotefuran colony became rapidly extinct within a month on August 16^th^ when none of adult bees and capped brood existed.

The fenitrothion colony (RUN3) shows a decrease of 13.3 percent in the number of adult bees within a day from July 21^st^ to July 22^nd^ in comparison with the initial number on July 21^st^. The decrease in the number of adult bees (1193; 13.3 percent) is somewhat more than the number of dead bees (865; 9.7 %) in the same interval. This suggests that most of dead bees died on the spot and some of them became lost. The fenitrothion colony (RUN3) shows a decrease of 33.3 percent in the number of adult bees and a decrease of 93.0 percent in the number of capped brood on August 16^th^ in comparison with the initial number on July 21^st^.

At the elapse of 26 days the decrement of adult bees is 352.81 bees/day (9173 bees/26 days) in the dinotefuran colony and 114.69 bees/day ((8943-5961) bees/26days) in the fenitrothion colony. It can be seen from this that dinotefuran caused a decrease in the number of the adult bees in the colony about three times faster than that of fenitrothion.

According to the recovery experiment from August 16^th^ when the dinotefuran colony became extinct, it was found that the fenitrothion colony began to recover from the effect of fenitrothion immediately after the discontinuance of its administration. The number of capped brood reached to the minimum (7% of the initial) at the stop of fenitrothion administration on August 16^th^ and it immediately began to increase. The number of adult bees in the fenitrothion colony reached to the minimum (60 *%* of the initial) on September 6^th^ after 21 days elapsed from August 16^th^ when pesticide-free sugar syrup was fed into the fenitrothion colony. These facts suggest that fenitrothion adversely affects the oviposition of the queen during administration of fenitrothion but the adverse effect becomes virtually absent in a short period. The delay of 21 days to the minimum number of adult bees from that of capped brood seems to be due to the period for capped brood group of minimum number to grow up into the adult bee group of minimum number. The fenitrothion colony increased in the numbers of adult bees and capped brood as rapidly as both control colonies after wintering. This means that organophosphates such as fenitrothion can hardly exert a long-term effect on a honeybee colony and the chronic toxicity can be neglected. Though the control colony of RUN4 was attacked by Asian giant hornets with some bees being killed, it is not affected by them very much.

### The interval number of newly emerging adult bees estimated from capped brood between two adjacent observational dates

Now, we will estimate the interval number of adult bees which are newly emerging from capped brood (pupae) between two adjacent observational dates, that is, an observational date and the next one under the following assumptions (1) to (5) while giving examples of the dinotefuran colony (RUN2) based on the experimental data in Table 3. All of the newly emerging adult bees during administration period are assumed to have taken the pesticide.

(1) The age distribution of the capped brood at an observation date is uniform between the first day when the cells of larvae are newly capped and the twelfth day when they eclose. (2) The number of adult bees that emerge from the pupae (capped brood) per day at a given day is one-twelfth of the number of the capped brood at the last observation date before the day. (3) The interval number of adult bees born between two successive observation dates is given by the product of one-twelfth of the number of the capped brood at the former observation date and the number of days from the former to the latter observation date. Here we will explain the procedure with examples: the interval number of newly emerging adult bees from capped brood can be obtained from the relation that (the number of capped brood/12)×(the number of days between two adjacent observational dates); that is, for the dinotefuran colony (RUN2), 9442/12[bees/day]×1[day]=786.83[bees]; 8834/12[bees/day]×5 [days]=3680.83[bees]; 4548/12[bees/day]×1[day]=379.00[bees];3891/12[bees/day]×6[days]= 1945.5[bees]; 1131/12[bees/day]×1[day]=94.25[bees]; and 840/12 [bees/day]×4[days]=280.00 [bees] between July 21^st^ of 2012 and July 22^nd^; July 22^nd^ and July 27^th^; July 27^th^ and July 28^th^; July 28^th^ and August 3^rd^; August 3^rd^ and August 4^th^; August 4^th^ and August 8^th^, respectively. (4) The procedure in (3) is applied even when the number of days between two successive observation dates is greater than 12. (5) The number of the capped brood at the time of the final pesticide administration or colony extinction is regarded as the number of adult bees having ingested the pesticide assuming that all the capped brood has already ingested the pesticide. Exceptionally, when the number of the capped brood at the colony extinction is zero, the number of newborn adult bees during the final interval is assumed to be equal to the number of the capped brood at the last observation before the final (extinction). For an example of the dinotefuran colony (RUN2), the number of capped brood on August 8th is 208 which have already taken the pesticide (dinotefuran). All the capped brood on August 8^th^ has emerged before August 16^th^ when the experiment of the dinotefuran colony (RUN2) has finished in this case.

The total number of newly emerging adult bees during the administration of pesticide can be obtained from the sum of the interval numbers of adult bees which are newly emerging from capped brood between two adjacent observational dates.

### The grand total number of honeybees during the administration period of pesticide

As the grand total number of honeybees during the administration period of pesticide is given by finding the sum of the total number of newly emerging adult bees during the administration period, the number of initial adult bees which have already existed at the start of experiment and the number of the capped brood at the end of the administration which have already taken a pesticide.

Here we will obtain the grand total number of honeybees from the start of experiment to August 16^th^ when the dinotefuran colony (RUN2) became extinct and the administration of fenitrothion was discontinued into the fenitrothion colony (RUN3).

For the dinotefuran colony (RUN2), the number of initial adult bees is 9173; the total number (sum of the interval numbers) of adult bees newborn between each two successive observation dates = (9442/12)(1) from July 21^st^ to 22^nd^ + (8834/12)(5) from July 22^nd^ to 27^th^ + (4548/12)(1) from July 27^th^ to 28^th^ + (3891/12)(6) from July 28^th^ to August 3^rd^ + (1131/12)(1) from August 3^rd^ to 4^th^ + (840/12)(4) from August 4^th^ to 8^th^ = 7166.4; and the number of newborn adult bees during the final interval from August 8^th^ to 16^th^, where they seem to have taken the pesticide (dinotefuran) before capped, is 208 that is the number of capped brood on August 8^th^, because capped brood was zero at the colony extinction on August 16^th^. That is, the grand total number of honeybees which have taken the pesticide in the dinotefuran colony (RUN2) is the sum (16547.4) of the number of the initial bees (9173), the total number of the newborn bees (7166.4) and the number of the final capped brood (208).

For the fenitrothion colony (RUN3), the number of initial adult bees is 8943; the total number (sum of the interval numbers) of adult bees newborn between each two successive observation dates = (8732/12)(1) from July 21^st^ to 22^nd^ + (8694/12)(5) from July 22^nd^ to 27^th^ + (6563/12)(1) from July 27^th^ to 28^th^ + (6389/12)(6) from July 28^th^ to August 3^rd^ + (3390/12)(1) from August 3^rd^ to 4^th^ + (2901/12)(4) from August 4^th^ to 8^th^ + (1352/12)(8) from August 8^th^ to 16^th^ = 10242.4; The number of the capped brood at the stop of administration of the pesticide (fenitrothion) on August 16^th^ was 607, all of which seemed to take the pesticide. That is, the grand total number of honeybees which took the pesticide in the fenitrothion colony (RUN3) is the sum (19792.4) of the number of the initial bees (18943), the total number of newborn bees (10242.4) and the number of the final capped brood (607).

### Intake of pesticide by a colony

The cumulative consumption of sugar syrup by each colony is shown in Figure 5. Table 4 shows the interval consumption of toxic sugar syrup with each pesticide ingested by the dinotefuran colony (RUN2) and the fenitrothion one (RUN3) during an interval between 2 successive observation dates and the cumulative total consumption of sugar syrup from July 21^st^ in 2012 to August 16^th^. The cumulative total consumption of sugar syrup by the dinotefuran colony is 776 g and that by the fenitrothion colony is 1707 g during the administration of a pesticide (dinotefuran or fenitrothion) from July 21^st^ to August 16^th^.

**Figure 5.**
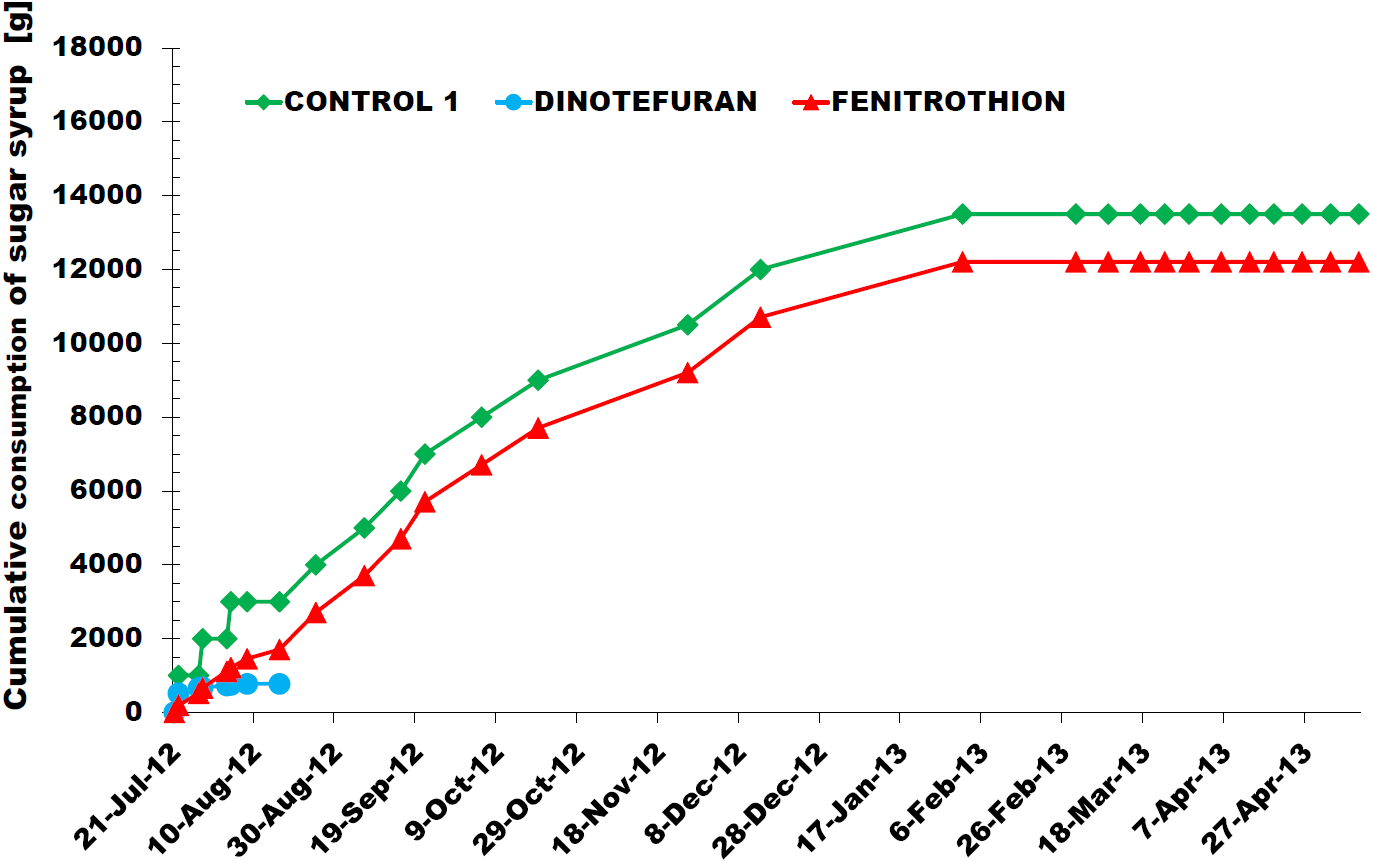
Cumulative consumption of sugar syrup by each colony. “Control 1”, “DINOTEFURAN” and “FENITROTHION” indicate the colonies supplied with sugar syrup containing no pesticide, dinotefuran and fenitrothion, respectively. These pesticides were administered into their target colonies from July 21^st^ to August 16 in 2012. Control 2 shows the same curve as Control 1. **Daily consumption of toxic sugar syrup with the pesticide by each colony** The daily consumption can be estimated by dividing a cumulative consumption of sugar syrup in a colony between adjacent observation dates by the number of days between them.

**Table 4.** Interval and cumulative consumptions of sugar syrup [g]

| Date | Elapsed days | RUN1 (Control 1)<br>without pesticide |  | RUN2 (Dinotefuran)<br>2 ppm |  | RUN3 (Fenitrothion)<br>10 ppm |  | RUN4 (Control 2)<br>without pesticide |  |
| --- | --- | --- | --- | --- | --- | --- | --- | --- | --- |
|  |  | Interval Consumption | Cumulative Consumption | Interval Consumption | Cumulative Consumption | Interval Consumption | Cumulative Consumption | Interval Consumption | Cumulative Consumption |
| 21-Jul-12 | 0 | 0 | 0 | 0 | 0 | 0 | 0 | 0 | 0 |
| 22-Jul-12 | 1 | 1000 | 1000 | 518 | 518 | 195 | 195 | 1000 | 1000 |
| 27-Jul-12 | 6 | 0 | 1000 | 145 | 663 | 324 | 519 | 0 | 1000 |
| 28-Jul-12 | 7 | 1000 | 2000 | 15 | 678 | 145 | 664 | 1000 | 2000 |
| 3-Aug-12 | 13 | 0 | 2000 | 50 | 728 | 452 | 1116 | 0 | 2000 |
| 4-Aug-12 | 14 | 1000 | 3000 | 20 | 748 | 100 | 1216 | 1000 | 3000 |
| 8-Aug-12 | 18 | 0 | 3000 | 28 | 776 | 239 | 1455 | 0 | 3000 |
| 16-Aug-12 | 26 | 0 | 3000 | 0 | 776 | 252 | 1707 | 0 | 3000 |
| 25-Aug-12 | 35 | 1000 | 4000 |  |  | 1000 | 2707 | 1000 | 4000 |
| 6-Sep-12 | 47 | 1000 | 5000 |  |  | 1000 | 3707 | 1000 | 5000 |
| 15-Sep-12 | 56 | 1000 | 6000 |  |  | 1000 | 4707 | 1000 | 6000 |
| 21-Sep-12 | 62 | 1000 | 7000 |  |  | 1000 | 5707 | 1000 | 7000 |
| 5-Oct-12 | 76 | 1000 | 8000 |  |  | 1000 | 6707 | 1000 | 8000 |
| 19-Oct-12 | 90 | 1000 | 9000 |  |  | 1000 | 7707 | 1000 | 9000 |
| 25-Nov-12 | 127 | 1500 | 10500 |  |  | 1500 | 9207 | 1500 | 10500 |
| 13-Dec-12 | 145 | 1500 | 12000 |  |  | 1500 | 10707 | 1500 | 12000 |
| 1-Feb-13 | 195 | 1500 | 13500 |  |  | 1500 | 12207 | 1500 | 13500 |
| 1-Mar-13 | 223 | 0 | 13500 |  |  | 0 | 12207 | 0 | 13500 |
| 9-Mar-13 | 231 | 0 | 13500 |  |  | 0 | 12207 | 0 | 13500 |
| 17-Mar-13 | 239 | 0 | 13500 |  |  | 0 | 12207 | 0 | 13500 |
| 23-Mar-13 | 245 | 0 | 13500 |  |  | 0 | 12207 | 0 | 13500 |
| 29-Mar-13 | 251 | 0 | 13500 |  |  | 0 | 12207 | 0 | 13500 |
| 6-Apr-13 | 259 | 0 | 13500 |  |  | 0 | 12207 | 0 | 13500 |
| 13-Apr-13 | 266 | 0 | 13500 |  |  | 0 | 12207 | 0 | 13500 |
| 19-Apr-13 | 272 | 0 | 13500 |  |  | 0 | 12207 | 0 | 13500 |
| 26-Apr-13 | 279 | 0 | 13500 |  |  | 0 | 12207 | 0 | 13500 |
| 3-May-13 | 309 | 0 | 13500 |  |  | 0 | 12207 | 0 | 13500 |
| 10-May-13 | 316 | 0 | 13500 |  |  | 0 | 12207 | 0 | 13500 |

Red figures denote toxic sugar syrup with the pesticide (dinotefuran or fenitrothion).

Assuming that the consumption of toxic sugar syrup per day is constant between 2 successive observation dates, the daily consumption can be estimated as shown in Table 5. It can be seen from Table 5 that the dinotefuran colony (RUN2) ingested about 67 percent (518g/776g) of the cumulative total comsumtion of toxic sugar syrup only within one day just after the first administration but the fenitrothion colony (RUN3) did no more than about 11 percent (195g/1707g). From another point of view, the initial daily consumption of toxic sugar syrup by the dinotefuran colony just after the first administration is about 2.7 times (518g/195g) as much as that by the fenitrothion. This difference may perhaps come from malodorous fenitrothion as opposed to odorless dinotefuran.

**Table 5.** Interval consumption of toxic sugar syrup from the start of administration (July 21^st^) to the finish (August 16^th^)

| Date | Elapsed days | RUN2 (Dinotefuran : 2ppm) |  | RUN3 (Fenitrothion : 10 ppm) |  | Note |
| --- | --- | --- | --- | --- | --- | --- |
|  |  | Interval consumption of toxic sugar syrup between 2 successive observation dates [g] | Daily consumption of toxic sugar syrup [g/day] | Interval consumption of toxic sugar syrup between 2 successive observation dates [g] | Daily consumption of toxic sugar syrup [g/day] |  |
| 21-Jul-12 | 0 | 0 | 0.00 | 0 | 0.00 | Observation date |
| 22-Jul-12 | 1 | 518 | 518.00 | 195 | 195.00 | Observation date |
| 23-Jul-12 | 2 |  | 29.00 |  | 64.80 |  |
| 24-Jul-12 | 3 |  | 29.00 |  | 64.80 |  |
| 25-Jul-12 | 4 |  | 29.00 |  | 64.80 |  |
| 26-Jul-12 | 5 |  | 29.00 |  | 64.80 |  |
| 27-Jul-12 | 6 | 145 | 29.00 | 324 | 64.80 | Observation date |
| 28-Jul-12 | 7 | 15 | 15.00 | 145 | 145.00 | Observation date |
| 29-Jul-12 | 8 |  | 8.33 |  | 75.33 |  |
| 30-Jul-12 | 9 |  | 8.33 |  | 75.33 |  |
| 31-Jul-12 | 10 |  | 8.33 |  | 75.33 |  |
| 1-Aug-12 | 11 |  | 8.33 |  | 75.33 |  |
| 2-Aug-12 | 12 |  | 8.33 |  | 75.33 |  |
| 3-Aug-12 | 13 | 50 | 8.33 | 452 | 75.33 | Observation date |
| 4-Aug-12 | 14 | 20 | 20.00 | 100 | 100.00 | Observation date |
| 5-Aug-12 | 15 |  | 7.00 |  | 59.75 |  |
| 6-Aug-12 | 16 |  | 7.00 |  | 59.75 |  |
| 7-Aug-12 | 17 |  | 7.00 |  | 59.75 |  |
| 8-Aug-12 | 18 | 28 | 7.00 | 239 | 59.75 | Observation date |
| 9-Aug-12 | 19 |  | 0.00 |  | 31.50 |  |
| 10-Aug-12 | 20 |  | 0.00 |  | 31.50 |  |
| 11-Aug-12 | 21 |  | 0.00 |  | 31.50 |  |
| 12-Aug-12 | 22 |  | 0.00 |  | 31.50 |  |
| 13-Aug-12 | 23 |  | 0.00 |  | 31.50 |  |
| 14-Aug-12 | 24 |  | 0.00 |  | 31.50 |  |
| 15-Aug-12 | 25 |  | 0.00 |  | 31.50 |  |
| 16-Aug-12 | 26 | 0 | 0.00 | 252 | 31.50 | Observation date |

(Note) Blue numbers are estimated from the consumption of sugar syrup measured at an observation date under the assumption that the consumption per day (consumption rate) is same between two successive observation dates from a certain observation date to the previous one: E.g., the consumption rate from July 23<sup>rd</sup> to 27<sup>th</sup> is obtained from dividing the consumption measured on July 27<sup>th</sup> (145g) by the interval (5 days) for RUN2.

The intake of a pesticide taken by each experimental colony is calculated from the cumulative total consumption of sugar syrup. As the concentration of dinotefuran in sugar syrup is 2 ppm and that of fenitrothion is 10 ppm, the cumulative total intake of dinotefuran becomes 1.552 mg and that of fenitrothion does 17.07 mg. Each cumulative total intake of pesticide means the intake of the pesticide (dinotefuran, fenitrothion) that honeybees of each colony removes from the feeder in the hive before August 16^th^. The cumulative total intake is the amount of the pesticide, some of which was ingested by honeybees and the others were stored as honey and bee bread in combs after honeybees converted toxic sugar syrup into toxic honey and/or toxic bee bread. That is, when toxic sugar syrup is stored as honey and/or bee bread, honeybees are inevitably affected by the pesticide through conversions. We cannot know the impact of the pesticide on honeybees when toxic sugar syrup is converted into honey and/or bee bread. We have to recognize that the cumulative total intake of the pesticide is not the true amount of the pesticide taken by honeybees but the apparent amount of the pesticide removed from the feeder to other places. Incidentally, we can relatively compare the cumulative total intakes under the same environmental conditions where the foraging activity seems to be about the same.

Here we will estimate the intake of pesticide per bee during the administration period of pesticide from dividing each cumulative total intake of dinotefuran or fenitrothion by the grand total number of honeybees during the administration period of pesticide. We can estimate the intake of pesticide per bee till the colony extinction of 93.8 ng/bee from dividing 1.552 mg by 16547.4 for the dinotefuran colony (RUN2) and that of 862.5 ng/bee from dividing 17.07 mg by 19792.4 for the fenitrothion colony (RUN3), respectively. Comparing the intake of dinotefuran per bee with the average LD_50_ for acute oral of a honeybee which is 20.9 ng/bee (7.6+23+32)/3), the ratio of the intake to the average LD_50_ is about 4.5. Similarly, the ratio of the intake of fenitrothion per bee to the LD_50_ for acute oral of a honeybee (200 ng/bee) is about 4.3. We can perceive that the intakes of the pesticides per bee are about 4.5 times higher than their LD_50_. This reason seems to be due to the amount of sugar syrup stored in combs, which can depend on the weather conditions, the blooming season of flowers and so on. Details will be discussed below.

## Discussion

### Differences in impact on a honeybee colony between dinotefuran and fenitrothion

Though we prepared toxic sugar syrup with both the concentration of dinotefuran and that of fenitrothion having one-fiftieth insecticidal activity to exterminate stinkbugs, we obtained very different results on the colony between the two pesticides: (1) The neonicotinoid dinotefuran colony (RUN2) became extinct after the elapse of 26 days from the administration of the pesticide but the organophosphate fenitrothion one (RUN3) did not and even could succeed in overwintering. (2) Dinotefuran can kill more than half the initial adult bees since immediately after the administration of the pesticide but fenitrothion can kill less than one-tenth of those with the same insecticidal activity for stinkbugs, while both pesticides seem to have almost the same impact adversely on the capped brood. (3) The initial consumption of toxic sugar syrup by the dinotefuran colony is two and a half times more than that by the fenitrothion colony. (4) The fenitrothion colony had a peak of the number of dead bees per day just after newly-prepared (fresh) toxic sugar syrup with the pesticide more clearly than the dinotefuran colony.

#### Why can dinotefuran kill more adult bees than fenitrothion?

We can find that from Figure 3 that adult bees in the dinotefuran colony steeply decreased in number just after the administration of dinotefuran and became extinct in a short period of time and those in the fenitrothion colony gradually decreased in number to about two-thirds of the initial at the discontinuation of fenitrothion administration (the extinction of the dinotefuran colony), reached to the minimum (three-fifths of the initial) afterwards and then began to increase in number during the recovery experiment with assuming almost the same aspect as the control colonies. On the other hand, we can find from Figure 4 that capped brood in both experimental colonies steeply decreased in number just after the administration of the pesticides and reached to the minimum (0 *%* for the dinotefuran colony of the initial; 7 % for the fenitrothion colony) at the extinction of the dinotefuran colony (the stop of pesticide administration) and then began to increase in number during the recovery experiment with assuming almost the same aspect as the control colonies.

It can be suggested that the insecticidal activity of fenitrothion is much weaker than that of dinotefuran despite their same insecticidal activity for a stinkbug as seen also from Figure 9, which shows the daily number of dead bees per adult bees (namely, mortality per day) expressed in value relative to that on July 21^st^. We can probably understand that the queen was severely affected adversely by the pesticides and her oviposition capacity was reduced when toxic sugar syrup with pesticide was given to the queen as toxic honey or toxic bee bread which was made by mixing pollen and toxic honey, and/or the brood were also affected adversely by the pesticides before being capped when toxic honey and toxic bee bread were given to them by house bees. Especially, bee bread seems to be given before the pesticides have lost their toxicity because of a short period of their storage in cells (Gillian, 1979; DeGrandi-Hoffman etal., 2013).

**Figure 6.**
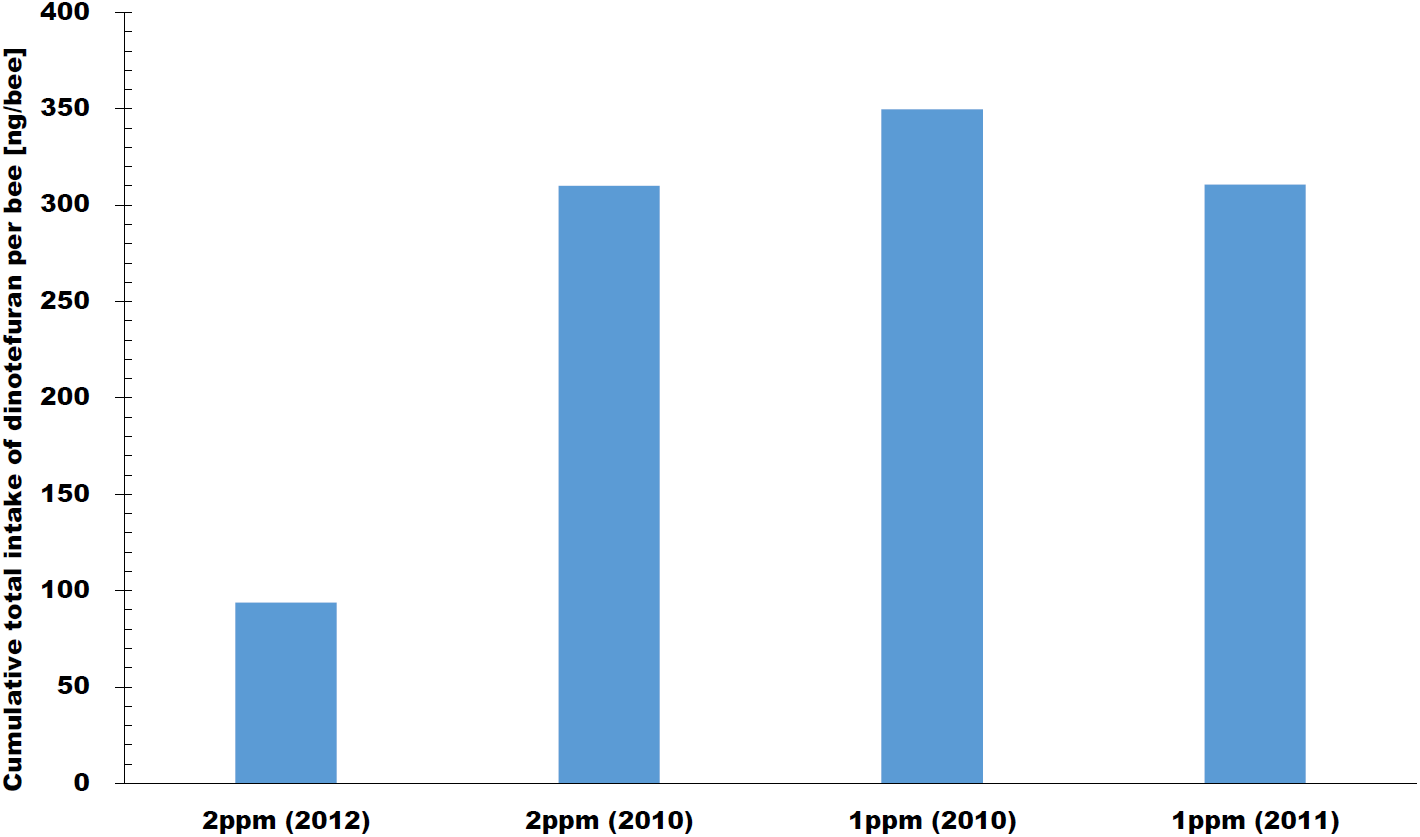
Estimated total intake of dinotefuran per bee till the colony extinction in this work and previous ones. We compare the estimated amount of dinotefuran that a honeybee takes till the colony extinction among three kinds of our field experiments which started at 2010, 2011 and 2012. Each concentration such as 2 ppm indicates the concentration of dinotefuran in sugar syrup fed to a colony. The number in the parenthesis indicates the year for each of our field experiments: The year of 2012 indicates this work, and the other years of 2010 and 2011 indicate our previous works which have been already reported by Yamada **et al.** (2012) and Yamada **et al.** (under submission), respectively.

**Figure 7.**
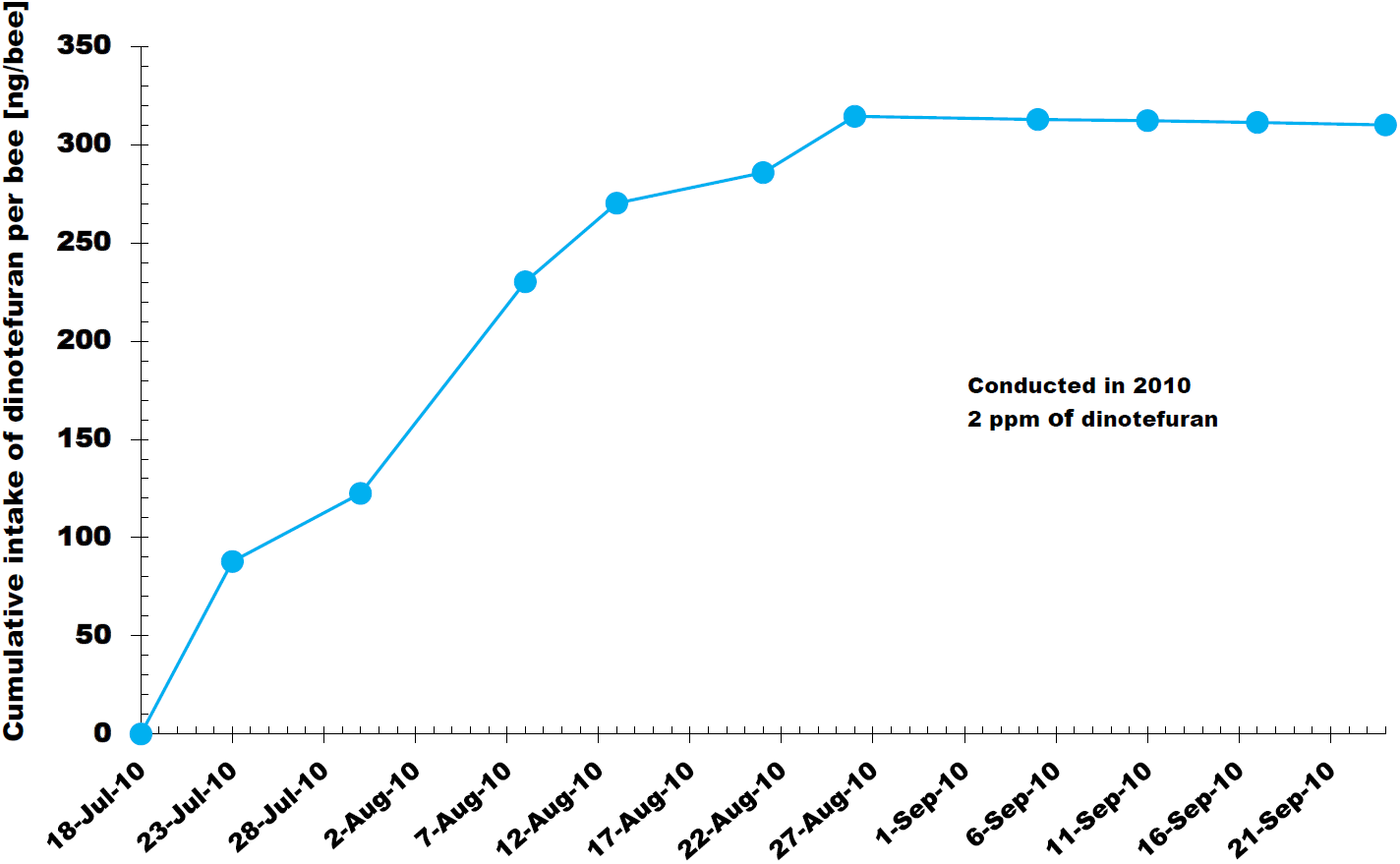
Cumulative intake of dinotefuran per bee in 2010. The cumulative intake of dinotefuran can be obtained by dividing a total of the intake by that of honeybees from the start of administration of dinotefuran till a certain observation date when the experiment was conducted in 2010 (Yamada *et al.*, 2012).

**Figure 8.**
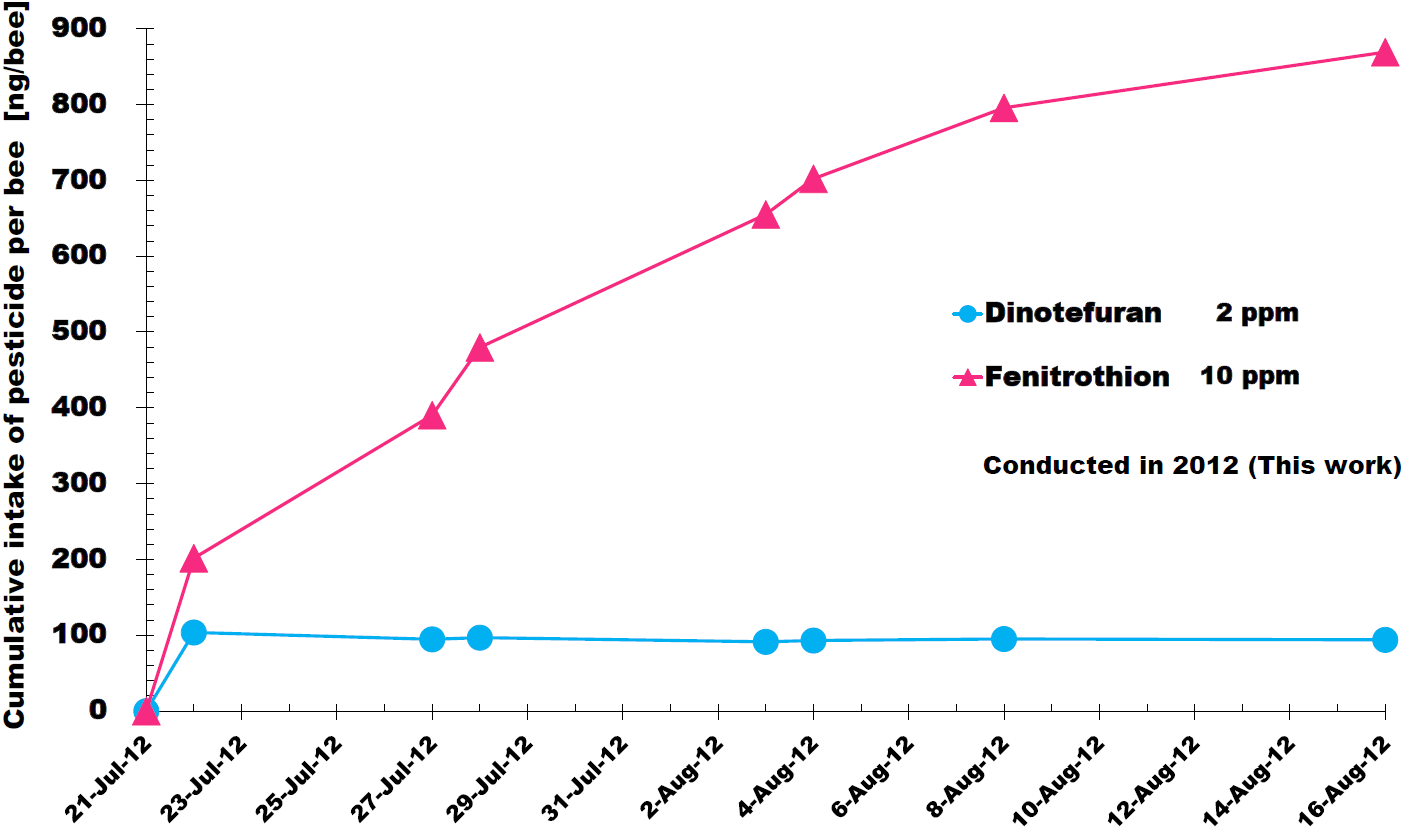
Cumulative intakes of dinotefuran and fenitrothion per bee in this work. These cumulative intakes can be obtained by the similar procedure to Figure 7.

**Figure 9.**
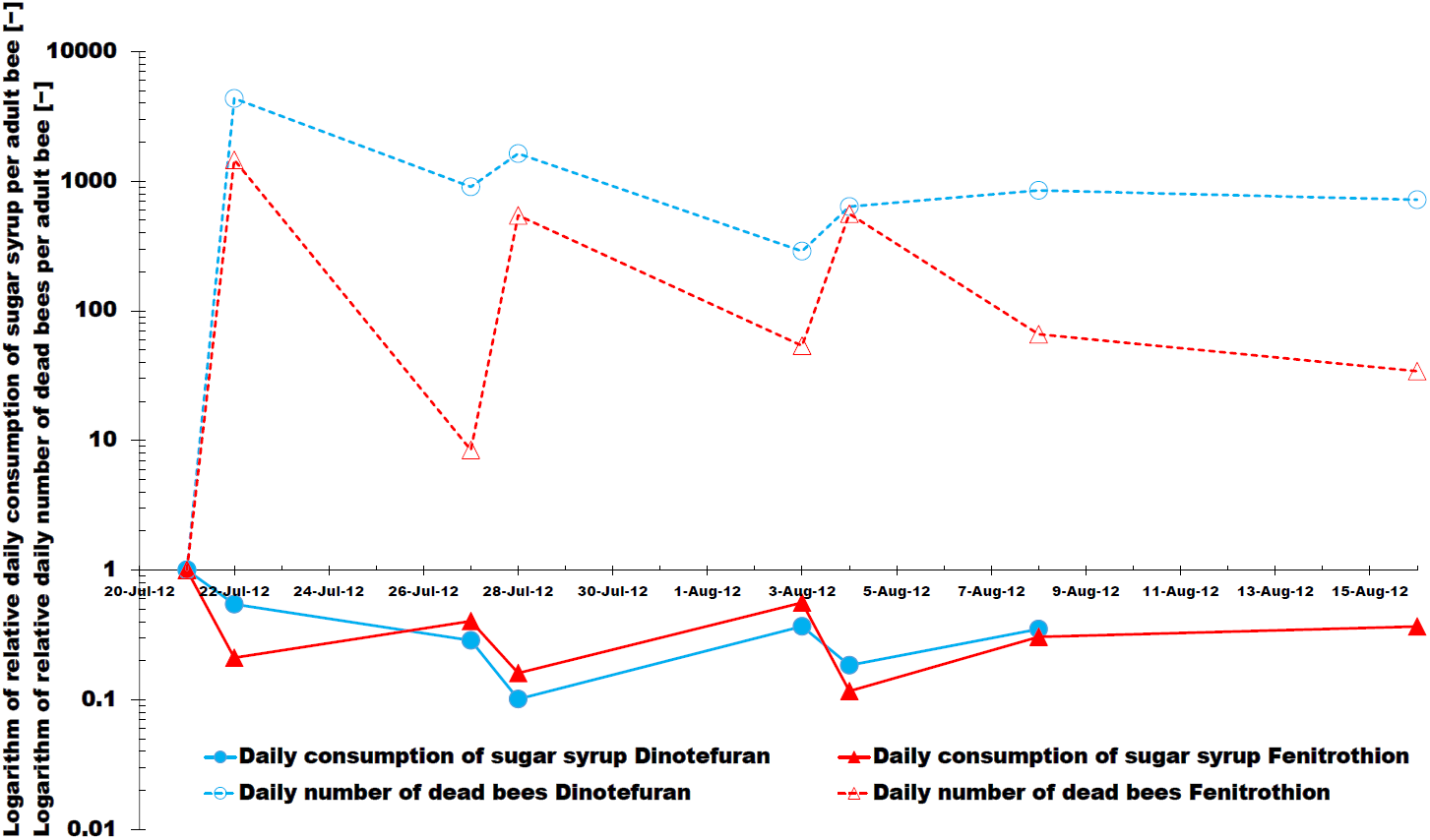
Daily consumption of sugar syrup per adult bee and daily number of dead bees per adult bee. The daily interval consumption of sugar syrup per adult bee [g/day/adult bee] is obtained from dividing the interval consumption of sugar syrup shown in Table 4 by the number of days in the interval between two adjacent observation dates and by the last number of adult bees before an observation date. The daily number of dead bees per adult bee (that is, mortality per day) [heads/day/adult bee] is obtained from dividing the interval number of dead bees shown in Table 2 by the number of days in the interval between two adjacent observation dates and by the last number of adult bees before an observation date. The relative values to a standard are shown in this figure. A standard of the daily consumption of sugar syrup per adult bee in assuming that each colony takes nontoxic sugar syrup is the average quantity of sugar syrup consumed by two control colonies for a day from July 21^st^ to July 22^nd^ as a substitute for the nontoxic quantity before the administration of the pesticide into each experimental colony because we have not measured the nontoxic quantity before the administration; 1000g/9647 heads for Control 1 (RUN1) and 1000g/9665 heads for Control-2 (RUN4) A standard of the daily number of dead bee per adult bee for each experimental colony before the pesticide administration is obtained from dividing the number of dead bees measured one on July 21^st^ (5 heads for the dinotefuran colony; 3 heads for the fenitrothion one) by the number of days from July 15^th^ to July 21^st^ (6 days) and by the number of adult bees on July 15^th^ (6878 heads for the dinotefuran colony (RUN2); 7565 heads for the fenitrothion one (RUN3)) Their common logarithmic values are plotted except when they become zero. We assumed that the daily consumption of nontoxic (pesticide-free) sugar syrup per adult bee on July 21^st^ before the administration of the pesticide seems to be almost the same as the average daily consumption of nontoxic sugar syrup per adult bee by the two control colonies (Control 1 and Control 2) from July 21^st^ to July 22^nd^, where the average value of two controls is 0.1036 g/day/adult bee; namely, 0.1037 g/day/adult bee in Contro1 1 and 0.1035 g/day/adult bee in Contro1 2. The dates in 2014 when the fresh pesticide instead of old one was administered are as follows: The first pesticide administration date: July 21^st^; the second date: July 27^th^ and the third date: August 3^rd^.

Incidentally, we will deduce a factor which causes the difference between dinotefuran and fenitrothion from the following hypothesis about neurotransmission: Supposing that the frequency and quantity of acetylcholine (ACh) differ among a brood (larva), an adult bee (worker) and a queen, those of the enzyme acetylcholinesterase (AChE) which generates in order to readily decompose ACh may also differ among them (Dewhurst *et al.*, 1970; Grzelak, *et al.*, 1970; Mohamad, 1982; van der Kloot, 1955). That is to say, an adult bee without peculiar behavior produces less ACh and less AChE than a brood with feeding behavior and a queen with ovipositional behavior.

Assuming that ACh which can activate non-specific cation conductance to directly excite neurons is produced more in a brood which has to aggressively inform a nurse bee that she needs her feed than in an adult bee, AChE in the brood becomes more than that in the adult bee. As the neonicotinoid dinotefuran acts as an agonist of the ACh receptor by binding to the postsynaptic nicotinic acetylcholine receptor and the nerve is continually stimulated by dinotefuran itself while AChE is not affected by it, dinotefuran act on the nervous system independently of the frequency and quantity of actual ACh. As a result, dinotefuran seems to exhibit similar toxicity for an adult bee to that for a brood.

On the other hand, as the organophosphate fenitrothion acts on the nervous system as inhibitor of AChE and continued transmission of ACh, fenitrothion strongly affects AChE. As a result, fenitrothion which can decompose AChE probably continue to stimulate the nervous system of a brood stronger than that of an adult bee though dinotefuran which is an acetylcholine mimic and cannot be influenced by AChE continues to strongly stimulate the nervous system of a brood similarly to that of an adult bee regardless of the frequency and quantity of AChE.

Now, we will consider the influence of these pesticides on the nervous system of a queen where ACh seems to generate when she oviposits. Considering that AChE which generates as ACh generates is decomposed by fenitrothion, ACh can continue to affect the nervous system of the queen similarly to a brood under the condition of little AChE and can reduce her oviposition activity. Dinotefuran mimicking ACh also continue to affect the nervous system of the queen, unaffected by AChE, and reduce her oviposition activity, as is the case with fenitrothion. That is, a queen exposed to fenitrothion seems to lay almost the same small number of eggs as dinotefuran.

#### Why can the dinotefuran colony consume toxic sugar syrup at the first administration more than the fenitrothion colony?

Figure 10 shows the consumption of toxic sugar syrup with 2 ppm dinotefuran taken by the colony during each interval between two observation dates and that of toxic sugar syrup with 10 ppm fenitrothion in this work. It can be seen from this figure that the dinotefuran colony takes an extremely larger quantity of toxic sugar syrup (about 2.7 times as much as) than the fenitrothion colony just after the first administration, when the numbers of adult bees and capped brood in each colony were on almost the same level, respectively, after the acclimatization period. This tendency can be seen in the daily consumption of toxic sugar syrup per adult bee in Figure 9 which shows the daily consumption of toxic sugar syrup per adult bee which is obtained by dividing daily consumption of toxic sugar syrup by the last number of adult bees before an observation date. These suggest that fenitrothion seems to be more repellent than dinotefuran as pointed by Kegley *et al.* (2014) about a slight repellent effect of organophosphates such as fenitrothion and GELS *et al.* (2002), Larson *et al.* (2013) and BASF (2014) about a non-repellent effect of neonicotinoids such as dinotefuran.

**Figure 10.**
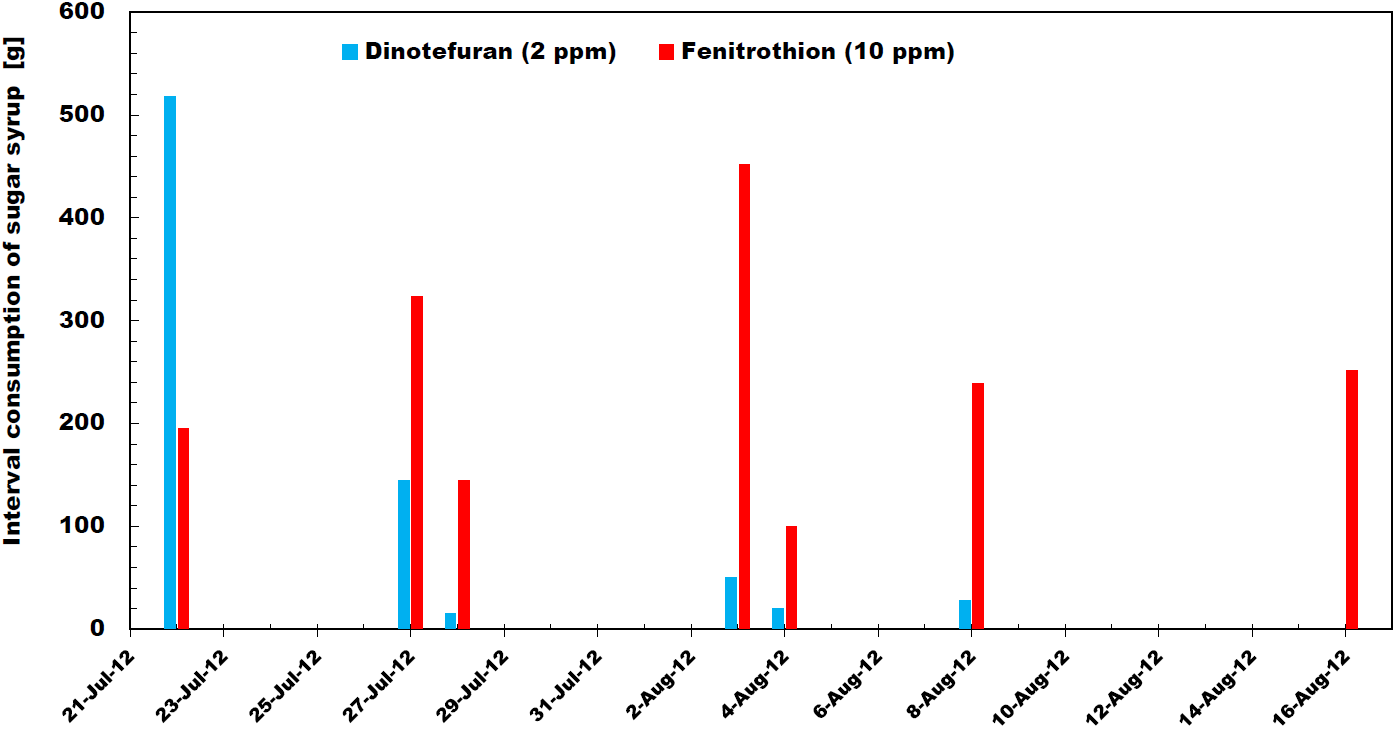
Interval intake of sugar syrup with the pesticide for each colony between two adjacent observation dates in this work. The interval intake of sugar syrup can be obtained by the amount of sugar syrup consumed by each colony between two adjacent observation dates: Ex. The interval intake on July 27^th^ is the amount of sugar syrup consumed from July 22^nd^ till 27^th^ in a colony.

#### Why can fresh toxic sugar syrup with fenitrothion kill more adult bees than older one?

As shown in Figure 2, daily dead bees in the fenitrothion colony rapidly increase in number just after feeding newly-prepared toxic sugar syrup with fenitrothion into the hive and afterwards begin to decrease in number every administration. On the other hand, the tendency is not clearly visible for those in the dinotefuran colony. The daily number of dead bees is obtained by dividing the interval number of dead bees by the number of days in the interval referring to Table 2. As the interval number of dead bees depends on the population to which they belong, we try to obtain the daily number of dead bees per adult bee which is obtained from dividing the daily number of dead bees by the population (the number of adult bees shown in Table 3) at the last observation before counting the dead bees which seem to have belonged there. The relative daily number of dead bees per adult bee is shown in Figure 9 after the conversion to a logarithmic scale, which is shown in value relative to that on July 21^st^ before the administration for each experimental colony (0.0001212 heads/day/adult bee on July 21^st^ for the dinotefuran colony and 0.00006609 heads/day/adult bee for the fenitrothion colony). From this figure we can find that the daily number of dead bees per adult bee for the fenitrothion colony shows the extremely clear tendency in rapid increase and that for the dinotefuran colony shows the slightly visible tendency. Noticeably the daily number of dead bees per adult bee for the fenitrothion colony is much lower than that for the dinotefuran colony.

Here we will examine the daily consumption of sugar syrup per adult bee. As the consumption of toxic sugar syrup by honeybees also depends on the population to which they belong, we try to obtain the daily consumption of toxic sugar syrup per adult bee by dividing the daily consumption of toxic sugar syrup shown in Table 5 by the population (number of adult bees shown in Table 3) at the last observation before counting the dead bees which seem to have belonged there. The daily consumption of toxic sugar syrup per adult bee is shown in Figure 9 after the conversion to a logarithmic scale, which is shown in value relative to the average of those by two control colonies for a day between July 21^st^ to 22^nd^ (0.1036 g/day/adult bee; namely, the average of 0.1037 g/day/adult bee in Contro1 1 and 0.1035 g/day/adult bee in Contro1 2. From this figure we can find that the daily consumption of toxic sugar syrup per adult bee for each experimental colony change with time as follows: At the elapse of a day after the first administration on July 21^st^ in 2012, the daily consumption of toxic sugar syrup per adult bee by the dinotefuran colony is much greater than that for the fenitrothion colony. After that, their relationship is reversed so that the daily consumption of toxic sugar syrup per adult bee by the fenitrothion colony is greater than that for the dinotefuran one on July 27^th^. Thereafter, the daily consumptions of toxic sugar syrup per adult bee for both experimental colonies similarly show the clear tendency in decrease just after feeding newly-prepared toxic sugar syrup and afterwards begin to gradually increase. This change in consumption of toxic sugar syrup after the second administration is quite contrary to that in dead bees.

Examining Figure 9 in more details, we can find that the daily number of dead bees per adult bee much more decreases after the first administration of fenitrothion than dinotefuran, and subsequently it turns to a much sharper increase after the second administration. These tendency recurs with attenuating the amplitude of vibration every administration. The daily number of dead bees per adult bee in the dinotefuran colony becomes almost constant keeping its peak after the third (final) administration and that in the fenitrothion colony begins to decrease after the final peak. The daily number of dead bees per adult bee keeps the level much higher in the dinotefuran colony than in fenitrothion colony and after the third administration the difference between the two colonies widens. These suggest that the insecticidal activity of fenitrothion can decrease with time much more rapidly than that of dinotefuran. It seems probable that easy decomposability and short-term persistence of fenitrothion (Pehkonen & Zhang, 2002) can cause the decrease in toxicity with time.

Here we will discuss in detail the daily consumption of toxic sugar syrup per adult bee shown in Figure 9. Just before the first administration, we did not measure the daily consumption of toxic sugar syrup per adult by each colony. If each of that just before the first administration is almost the same as the average of those by the control colonies between July 21^st^ and July 22^nd^, it is roughly 0.1036 g/day/adult bee (0.1037 g/day/adult bee for RUN1 (Control 1); 0.1035 g/day/adult bee for RUN4 (Control 2)) from Tables 3 and 4. Permitting the above assumption, the daily consumptions of toxic sugar syrup per adult bee for both experimental colonies rapidly decrease just after every administration.

The above rapid decrease in the intake of toxic sugar syrup just after every administration seems to be due to the following reasons: Firstly, the rapid decrease can be caused by repellency due to volatile constituents (Debboun *et al.*, 2006; Jacob John *et al.*, 2007) included in the pesticide consisting of not only the active ingredient but also inactive ones such as adjuvants and additives because the fresh pesticide usually includes more volatile constituents than the old one. Secondly, the disturbance of each colony due to our observation in the hive causes a reduction in foraging activity and therefore that honeybees seem to directly ingest toxic sugar syrup more, which cannot be stored in combs, than nontoxic nectar in fields gives rise to massive death of honeybees by a smaller amount of toxic sugar than in each interval.

Except for the first interval after the first dinotefuran administration, the daily consumption of toxic sugar syrup per bee by each experimental colony gradually increases with time, reaches its peak before the next administration and decreases just after the next administration, afterwards repeating the similar tendency. After the third (final) administration, it gradually increases with time. In the first interval the daily consumption of toxic sugar syrup per adult bee by the dinotefuran colony gradually decreases with time conversely.

The gradual decrease in the daily consumption of toxic sugar syrup in the first interval by the dinotefuran colony seems to due to the following reasons: Firstly, the dinotefuran colony has taken a great amount of toxic sugar syrup and stores some in combs after conversion into toxic honey after the first administration. We can infer that as the stored toxic sugar syrup (honey) continues to be consumed by the dinotefuran colony after the first administration, the daily consumption slightly decrease with time after the first administration. Secondly, the dinotefuran colony can be enfeebled by a great deal of the intake of toxic sugar syrup with dinotefuran just after the first administration and therefore honeybees can lose their appetite.

The gradual increase in the daily consumption of toxic sugar syrup by each experimental colony in each interval other than that by the dinotefuran colony in the first interval seems to be the following reasons: Firstly, a decrease in volatile constituents included in the pesticide with time causes an increase in the consumption of toxic sugar syrup which is considerably stored in combs, considering also the facts that mosquitoes are able to ignore the smell of the insect repellent within a few hours of being exposed to it (Stanczyk *et al.*, 2013) and organophosphates induce a phenomenon that was first attributed to repellency for foraging bees (Belzunces *et al.*, 2012). Secondly, capped brood which could take less toxic sugar syrup newly eclose in each interval in the colony and they consume toxic sugar syrup because they are more active than honeybees which already have ingested the pesticide.

Despite of the almost the same level of the daily consumptions of both pesticides, the much higher level of the daily number of dead bees in the dinotefuran colony than the fenitorothion colony means that dinotefuran seems to be higher toxic for a honeybee than fenitrothion under the same insecticidal activity for a stinkbug.

Incidentally, we should consider that these consumption of toxic sugar syrup and number of dead bee per adult bee can contain some margin of error when the population to which the adult bees belong is small.

#### Why can the fenitrothion colony succeed in overwintering?

We can find the following big difference between the neonicotinoid dinotefuran and the organophosphate fenitrothion with the same insecticidal activity for stinkbugs as each other: The dinotefuran colony became rapidly extinct within a month, while the fenitrothion colony even succeeded in overwintering instead of having taken a substantial amount of toxic sugar syrup. It seems probable that easy decomposability and short-term persistence of fenitrothion can lead to succeed in overwintering and recovering for the fenitrothion colony.

Here we will examine our previous works on the recovery experiments, strictly speaking, though they were conducted under different experimental conditions from this work: The neonicotinoids dinotefuran and clothianidin colonies had never been able to recover even after both the pesticides having one-tenth insecticidal activity to exterminate stinkbugs were administered only once and soon we converted from toxic foods (sugar syrup, pollen paste) to pesticide-free foods. That is probably attributed to the long-term persistence of neonicotinoids as reported by Yamada *et al.* (2012). In addition, we have the fact that the dinotefuran colony, where a low concentration of dinotefuran (0.565 ppm) was administered through pollen paste into which nontoxic pollen was kneaded with toxic sugar syrup having one-hundredth insecticidal activity to exterminate stinkbugs, failed in overwintering at the intake of dinotefuran of about 61 ng/bee, as reported by Yamada *et al.* (under submission) though it looked vigorous before winter. It can be deduced from these findings that neonicotinoids can cause not only CCD but also a failure in overwintering.

### Difference in the survival period of the dinotefuran colony between this work and previous work (Yamada *et al.*, 2012)

The dinotefuran colony in this work led to the much more rapid extinction (26 days) than that (61 days) in previous work reported by Yamada *et al.* (2012) under the same concentration. We will discuss from how such an inconsistency could arise.

Table 6 shows the cumulative total intakes of dinotefuran per bee till the colony extinction in this work, in comparison with those in our previous works experimented in 2010 (Yamada *et al.*, 2012) and in 2011 (Yamada *et al.*, under submission). Figure 6 shows the comparison of the estimated cumulative total intakes of dinotefuran per bee till the extinction of colony among our field-experimental results. We can find from Figure 6 and Table 6 that there is a big difference of the cumulative total intakes of pesticide per bee between this work and previous ones as follows: In this work conducted at the concentration of 2 ppm, we observed that more than half the initial number of honeybees died within a day after the first administration and the colony became extinct after the elapse of 26 days while a honeybee was estimated to take dinotefuran of 93.8 ng/bee. In previous work conducted at the concentration of 2 ppm in 2010 (Yamada *et al.*, 2012), a number of dead bees occurred only in the early period after the start of administration but they almost never occurred afterwards and the colony became extinct after the elapse of 61 days while a honeybee was estimated to take dinotefuran of 310.0 ng/bee. In other previous works conducted at the concentration of 1 ppm in 2010 and 2011, dead bees almost never occurred after the administration and the colony became extinct after the elapses of 84 days in 2010 (Yamada *et al.*, 2012) and 104 days in 2011 (Yamada *et al.*, under submission) while a honeybee was estimated to take dinotefuran of 349.8 ng/bee and 310.7 ng/bee in 2011. The colony extinction in this work seems to be chiefly triggered by a massive death due to acute toxicity, while the extinction in previous works seems to be caused by chronic toxicity with assuming an aspect of CCD.

**Table 6.** Intake of the pesticide per bee till the colony extinction (during administration) The fenitrothion colony (RUN3) in this work did not become extinct, the intake per bee was estimated using the cumulative number of honeybees and the cumulative total intake of fenitrothion taken by honeybees from the start of the pesticide administration on July 21^st^ in 2012 to the finish on August 16^th^ in 2012 when the dinotefuran colony (RUN2) in this work became extinct.

|  | 2012 (DF-2 ppm) | 2011 (DF-1 ppm)<sup>1)</sup> | 2010 (DF-1 ppm)<sup>2)</sup> | 2010 (DF-2 ppm)<sup>2)</sup> | 2012 (FT-10 ppm) |
| --- | --- | --- | --- | --- | --- |
| **Pesticide** | Dinotefuran | Dinotefuran | Dinotefuran | Dinotefuran | Fenitrothion |
| **Concentration of pesticide in vehicle** | 2 ppm | 1 ppm | 1 ppm | 2 ppm | 10 ppm |
| **Dilution factor against a concentration to exterminate stinkbugs** | a fiftieth part to exterminate stinkbugs | a hundredth part to exterminate stinkbugs | a hundredth part to exterminate stinkbugs | a fiftieth part to exterminate stinkbugs | a fiftieth part to exterminate stinkbugs |
| **Vehicle to administer the pesticide** | sugar syrup | sugar syrup | both sugar syrup and pollen paste | both sugar syrup and pollen paste | sugar syrup |
| **Notation** | DF-Middle | DF-Low | DF-Low | DF-Middle | FT-Middle |
| **Intake of the pesticide per bee till extinction [ng/bee]** | 93.8 | 310.7 | 349.8 | 310.0 | 862.5 |
| **Period to estimate the intake of pesticide** | From start to colony extinction | From start to colony extinction | From start to colony extinction | From start to colony extinction | From start to stop of pesticide administration |

1) Yamada *et al.*, under submission to Journal Apicultural Research
2) Yamada *et al.* (2012)

#### Why did the dinotefuran colony in this work become extinct by assuming an aspect of acute toxicity?

Now we will deduce the reason why the dinotefuran colony in this work became extinct after surviving for 26 days probably due to acute toxicity earlier than that in our previous work (Yamada *et al.*, 2012) had done after surviving for 61 days probably due to chronic toxicity under almost the same concentration of dinotefuran. In the field experiment of an actual apiary all of toxic sugar syrup with dinotefuran that is administered is not taken instantly, but it is stored as honey and the excipient of bee bread after the toxic sugar syrup was mixed by nectar or pollen without pesticides gathered from fields and the toxicity was attenuated. Considering that the amount and pesticide-concentration of stored toxic sugar syrup depend on the weather or the blooming season (Tesfay, 2007; Gebremedhn *et al.*, 2014), we here investigate them near the experimental site in Noto District where we conducted the experimental in our apiary in Ishikawa Prefecture, Japan. We cannot find the difference in blooming season between previous and this works because our previous pesticide-administration experiment in 2010 reported in Yamada *et al.* (2012) started on July 18^th^ and this one started on July 21^st^. Then we carefully investigate the weather for the initial period after toxic sugar syrup with dinotefuran started to be administered into a honeybee colony because the initial intake of the pesticide (dinotefuran) seems to most affect a honeybee colony.

Here we will examine the changes in maximum atmospheric temperatures of the days for about a month from the middle of July to the beginning of August in both 2010 and 2012 in Noto District near the experimental site (The Japan Weather Association). Comparing the changes in maximum atmospheric temperatures of the days for a month between in 2010 (Yamada *et al.*, 2012) and 2012 (this work), we can find that there was a significant difference between them for a week around the start of experiment. Examining a maximum atmospheric temperature of each day from three days before the start of experiment to three days after it, we can find the fact that the maximum, the minimum and the average among them are 34, 27 and 31.5 °C in 2010; and 30, 27 and 28.3 °C in 2012, respectively. Incidentally, the maximum, the minimum and the average of atmospheric temperatures for two week after the start of experiment were 34, 31 and 32.4 °C in 2010; and 36, 27 and 32.3 °C in 2012, respectively. We can find the fact that the temperatures just after the start of the experiment in 2012 (28.3 °C of the average) are lower than those in 2010 (31.5 °C of the average). The difference in temperature change between the two will discussed below:

Roughly speaking, the foraging activity (flight intensity) of honeybees tend to increase with temperature (Tesfay, 2007; Gebremedhn *et al.*, 2014). According to the number of honeybees visiting sunflower inflorescences during peak flowering when atmospheric temperature ranges from about 25°C to 35°C, it has been clarified that the foraging activity of honeybees increases sharply from about 25 °C to about 30°C, then it takes a maximum value at about 30 °C and then the maximum value is maintained till about 32°C, but after that it begins to decrease (Tesfay, 2007). Judging from the findings obtained by Tesfay (2007) and the temperature changes in our experimental site (Noto District in Japan), the foraging activity seems to remain high because the maximum temperatures were ranging from 31°C to 34 °C in 2010, but it seems to be fairly low for a few days just after the first administration of pesticide (dinotefuran) in 2012 in this work. Besides, the wide fluctuation of the temperatures from 27 °C to 36 °C in 2012 which take sometimes a value lower than 30°C or higher than 35°C can lead to a further decrease in foraging activity.

When the foraging activity is low, it will be generally accepted that honeybees bring less foods (nectar and pollen) from fields. From the above, it can be inferred that the dinotefuran colony in this work where the experiment was performed in 2012 brought less pesticide-free foods from fields, where we prepared a pesticide-free watering place and flower fields in our apiary, to the hive than the colony in our previous work where the experiment was performed in 2010 (Yamada *et al.*, 2012). It seems that some amount of foods consumed by honeybees is ingested by honeybees and the rest is stored in combs. It will be generally accepted that honeybees seem to prefer natural foods (nectar and pollen) to artificial foods (sugar syrup and pollen substitute) and they prefer nontoxic foods to toxic foods. From the above, we can infer that honeybees ingested foods in which a ratio of natural and nontoxic foods from our apiary to artificial toxic foods is higher in previous work in 2010 than in this work in 2012 and the intake of dinotefuran from ingested foods is less in our previous work in 2010 than in this work in 2012.

Meanwhile, we may infer that foods (honey and bee bread) stored in the hive for the colony in this work (conducted in 2012) becomes less than that in previous work (conducted in 2010) and the concentration of pesticide (dinotefuran) in stored foods in this work becomes higher than that in our previous work.

Therefore, honeybees actually ingested more pesticide (dinotefuran) and the colony became extinct in a shorter period of time after the first administration of pesticide assuming an aspect of acute toxicity in this work than in our previous work which had assumed an aspect of CCD, while the intake of dinotefuran per bee in this work was apparently less than that in our previous work

On the other hand, we have previously deduced that the main reason for few differences in the intake of dinotefuran per bee between the experimental results in 2010 and those in 2011 can come from few difference of change in atmospheric temperature between the two as reported previously (Yamada *et al.*, 2o12; Yamada *et al.*, under submission).

The difference in atmospheric temperature changes may probably cause the difference in the survival period of a colony as it was described above that the colony in this work became extinct earlier than that in our previous work. Here we should perceive in an experimental apiary that all of the amount of pesticide administered into a colony through food is not instantly taken by honeybees and some amount of the pesticide can be stored in combs after mixed with foods imported from fields where pesticides may or may not exist. In order to obtain the amount of the pesticide stored in the hive (combs), it may be necessary to accurately determine the amount of honey and bee bread in each comb and the concentration of the pesticide in them in every observation but it approaches the impossible. In this work we have relinquished their measurements.

#### Why is the intake of dinotefuran in this work less than that in our previous ones?

We have discussed above the difference in the survival period of the dinotefuran colony between this work and previous work (Yamada *et al.*, 2012). Here we will discuss the reason why the intake of dinotefuran per bee till the extinction of colony in this work (93.8 ng/bee) is less than that in our previous work (310 ng/bee) under almost the same concentration of dinotefuran as shown in Table 6 and Figure 6. As mentioned above, the lower atmospheric temperature in 2012 (this work) assumed to have led to the more substantial intake of dinotefuran in 2012 (this work) with the less toxic foods (honey, pollen) stored in combs than that in 2010 (Yamada *et al.*, 2012). Figure 7 and Figure 8 show the cumulative intake of dinotefuran taken by a honeybee till a certain observation date in our previous work conducted in 2010 and that in this work conducted in 2012, respectively, which is obtained from dividing the cumulative intakes of dinotefuran and fenitrothion taken by a colony (honeybees) till a certain observation date by a cumulative number of honeybees, which is given by the sum of both the initial number of adult bees at the start of experiment and the number of newborn bees till a certain observation date. Where the cumulative intake of fenitrothion per bee in this work is also shown in Figure 8 as a reference. Comparing these curves of dinotefuran between in 2010 and in 2012, we can find a difference between the two that the cumulative intake of dinotefuran in 2012 (this work) rapidly increases at the start of the experiment but that in 2010 (previous work) gradually increases. This may sustain the presumption mentioned that lower atmospheric temperatures leading to lower foraging activity cause more substantial intake of toxic food (sugar syrup, pollen paste) fed into a hive with less storing the food.

#### Can the LD_50_ assess the impact of a pesticide sprayed in fields on a honeybee colony in an apiary?

The LD_50_ is well-known as an indicator for acute toxicity of a pesticide. The LD_50_ for honeybees is defined by the amount of a pesticide which are individually forced to take and kills half of the honeybees within a limited time. The various values of the LD_50_ for fenitrothion have been reported by US-EPA (1995) (20 ng/bee for contact; 380 ng/bee for contact), Wang et al. (2012) (30 - 40 ng/bee for contact), Takeuchi et al. (1980) (130 ng/bee for contact), Okada and Hoshiba (1970) (30 ng/bee for contact), NUFARMNZ (2012) (18 ng/bee), University of Hertfordshire (2013) (160 ng/bee for contact) and Sanford (2003) (176 ng/bee for contact) and WHO (2010) (200 ng/bee for acute oral; 160 ng/bee for acute contact). The various values for dinotefuran also have been reported by US-EPA (2004) (23 ng/bee for acute oral; 47 ng/bee for contact; 32 ng/bee for acute oral; 61 ng/bee for contact; 7.6 ng/bee for acute oral; 24 ng/bee for contact), Iwasa et al. (2004) (75 ng/bee for contact) and Durkin (2009) (47 ng/bee for acute contact).

The LD_50_ is measured in the laboratory under controlled conditions, but in an actual apiary such as this field experiment site, there are many uncontrollable factors such as the behavior of a honeybee, environmental conditions, the weather, etc. Uncontrollable factors of environmental conditions and the weather can be cancelled to a certain degree by control experiment.

Judging from these LD_50_, the intake of the pesticide per bee as shown in our works are so high that the colony should be naturally expected to become extinct instantly but actually it has not done within a few days. Especially it is not understandable why the fenitrothion colony (RUN3) could even succeed in overwintering. One of possible causes is that the administration is not compulsory in the field experiment. The second is the stored toxic sugar syrup in cells on combs, which was diluted by pesticide-free honey from fields. This will be applicable to the dinotefuran colony (RUN2) because it continued to survive for 26 days while the cumulative total pesticide intake is enough to exterminate the colony within a few days.

In field conditions, a honey bee is free to go wherever she wants and take food whatever she wants, then she can selectively take food from fields if she prefer food in fields, which is unknown whether it is toxic or nontoxic, to toxic food with a pesticide administered. At a concentration of 2 ppm of dinotefuran in sugar syrup in this work, honeybees seem to be alive for a little while after the intake of the pesticide. While they are alive, they can convert toxic sugar syrup that they have taken from a feeder into toxic honey through a few honeybees and can temporally store it in combs.

Toxic honey can be mixed with honey made from nectar in fields in a cell or toxic sugar syrup can be mixed with nectar gathered from fields in honeybees’ bodies. Through a series of these processes, the toxicity of stored honey can be diluted. Pollen is kneaded with toxic honey to make bee bread and is stored in combs. In this work nectar and pollen from fields seems to be nontoxic because we have regulated our apiary to be pesticide-free though there is a slight possibility that they collect nectar and pollen from fields other than our apiary where pesticides are not controlled. The foods stored (honey, bee bread) are consumed by adult bees, brood and a queen.

The food containing neonicotinoids such as dinotefuran continue to adversely affect a honeybee colony for a long-term period of time but the food containing organophosphates cannot continue to affect a colony over a prolonged period because organophosphates such as fenitrothion can be decomposed easily and can be detoxicicated as shown above. This difference in persistence between organophosphates such as fenitrothion and neonicotinoids such as dinotefuran leads to a difference between success and failure in overwintering based on the fact that the fenitrothion colony in this work succeeded in overwintering but the dinotefuran colony in previous works (Yamada *et al.*, under submission) failed in overwintering though it looked vigorous before winter.

Besides the reasons mentioned above why the LD_50_ cannot assess the impact of a pesticide sprayed in fields on a honeybee colony in an apiary, we have to consider that the LD_50_ cannot give toxicological evaluations for a colony of honeybees which are eusocial insects because it can be used only to assess an individual living creature. We strongly desire a new indicator to assess chronic toxicity for a honeybee colony instead of the LD_50_.

#### How could CCD possibly be caused by a pesticide in an actual apiary?

It is defined as CCD that a honeybee colony exhibit all the following symptoms; a colony's worker bee population is suddenly lost with very few dead bees found near the colony; the queen and brood are remained; and the colonies had relatively abundant honey and pollen reserves; finally the colony cannot sustain itself without worker bees and would eventually die.

Now we will consider some plausible processes where a honeybee colony can assume an aspect of CCD by a pesticide in an actual apiary based on the fact that a honeybee is a eusocial insect.

When a pesticide is sprayed in fields, many foraging bees which are directly exposed to its high toxicity are instantly killed on the spot due to acute toxicity and the colony becomes short of foraging bees. Some house bees are recruited as foraging bees and then the caretakers of the brood become shorthanded in the colony. The queen lays less eggs. The colony dwindles away, become weakened and more susceptible to attack by pests and pathogens. Finally the colony cannot sustain itself and it collapses or escapes from the hive. In this case, CCD is hard to occur.

Here we will estimate how high the concentration of a pesticide causes an instant death of foraging bees in fields and makes foraging bees unable to return to their hive. A foraging bee has a honey stomach in which she can store 18 mg - 77 mg of nectar (Cooper *et al.*, 1985) and can carry about 40 mg of nectar (Yadav, 2003). Then the consumption of nectar per flight is about 13 mg under the assumption that the consumption of a foraging bee can be an approximate equivalent of the consumption of a drone (Burgett, 1973). When the concentration of a pesticide is x ppm, a foraging bee may carry 40x ng of a pesticide per flight and may take 13x ng of a pesticide during flight. Here a pesticide seems to act as a contact toxicity in case of being stored in the honey stomach of a foraging bee and being taken by her during transport. Now, taking dinotefuran as an example of a pesticide and assuming that the LD_50_ of dinotefuran is about 23 ng/bee for oral or about 61 ng/bee for contact (US-EPA, 2004) and most of foraging bees may die instantly on the spot at about twice the intake of a pesticide as much as the LD_50_, the thresholds of the pesticide concentration is about 3 ppm in honey stomach and about 3.5 ppm in her ingestion. That is, most of foraging bees which visit the field contaminated by dinotefuran of 3 ppm or more can probably be killed outright.

When the toxicity or concentration of the pesticide is not so high, many foraging bees indirectly exposed to the toxicity by which they cannot be killed on the spot and can bring toxic water, toxic foods (pollen, nectar) back to their hive. House bees directly consume some of them or store the other foods in combs after the toxic foods are diluted with nontoxic foods foraged from other uncontaminated fields. Some of honeybees exposed to the pesticide in the hive are killed in a short time due to acute toxicity or become weakened and get lost in fields depending on the amount of the pesticide taken by them because the stored ones continue to affect the colony adversely for a long period of time due to chronic if the pesticide is persistent. In this case, CCD can occur.

When most of foraging bees are not directly exposed to the pesticide and the toxicity of water, pollen and nectar in fields where the pesticide is sprayed is weakened by dilution with rainwater if the pesticide is water-soluble (systemic) and/or degradation due to sunlight. Foraging bees bring the weakened ones back to their hive and honeybees in the colony store some of foods in combs after the toxic foods are diluted with nontoxic foods foraged from other uncontaminated fields and directly consume others. Honeybees become weakened and get lost in fields due to chronic toxicity. The amount of toxic foods stored in combs depends on the foraging activity which is strongly influenced by environmental conditions such as weather and blooming conditions as can be seen from the difference in pesticide intake between this work and previous work (Yamada *et al,* 2012) under almost the same experimental conditions. Moreover, toxic water near fields contaminated with the pesticide also continue to adversely affect the colony while the toxicity is diluted with rainwater if the pesticide is persistent and is highly toxic. The stored foods and toxic water in fields continue to adversely affect the colony for a long period of time chronically if the pesticide is persistent. In this case, CCD can occur.

Even if the toxicity is too low to cause CCD during an active period of honeybees, it can cause a failure in overwintering due to chronic toxicity even when the colony looks vigorous before winter if the pesticide is persistent, because honeybees continue to ingest only the foods which are stored before winter and they live in winter several times as long as in active seasons.

We can infer that the disasters to a honeybee colony such as a CCD, a wintering loss and a massive death seems to be caused by the synergy effects due to a combination of the characteristics of a neonicotinoid pesticide such as long-term persistence, systemic property and high toxicity: The long term persistence permits the pesticide to long maintain its toxicity and to widely diffuse by dissolving in water in fields and being stored in a beehive as toxic foods under the natural environment; the systemic property permit it to easily dissolve in water and to be of wide distribution over the whole plant; and the high toxicity permits it to prolong its toxicity in a long period of time even after it is diluted by large quantities of rain water etc. On other hand, an organophosphate seems hard to cause such disasters except of a massive death just after being sprayed because it is probably much less persistent and lower toxic than a neonicotinoid

## Conclusion

From the field experiment from the end of June in 2012 to the middle of May in 2013, we confirm that dinotefran has much longer persistence on the honeybee colony in the field in comparison with fenitrothion. Although the concentrations of dinotefran and fenitrothion were adjusted to affect an individual bee with the similar level in terms of the LD_50_, there were clear differences between the colonies for which the different pesticides were administered: The dinotefran colony has become extinct within a month while the fenitrothion colony has succeeded in overwintering after the exposure.

Our results enlighten the persistent effects of pesticides in the field that cannot be estimated only from the LD_50_, i.e. acute toxicity for an individual that measured under laboratory conditions. The fenitrothion colony is estimated to have taken an enough amount of the pesticide to be extinct from the viewpoint of the short-term effects: During the administration, a bee in the fenitrothion colony is estimated to have taken 862.5ng of fenitrothion that is about 4.5 times larger than the LD_50_. The ratio of intake per bee to the LD_50_ of fenitrothion is comparable with that of dinotefran. Accordingly, the fenitrothion colony should be extinct at almost the same time as the dinotefran one if the LD_50_ can precisely evaluate the influence of the all kind of pesticides. Making an assessment of persistence of pesticides is urgent for the precise evaluation of the persistent toxicity to the wild animals and insects. To make an assessment, we need to pay more attention to complicated phenomenon itself in the natural environment that are often overlooked in the experiments in laboratory.

In addition, we found that dinotefuran and fenitrothion have shown the different impacts on the adult bees. Dinotefuran caused a decrease in the number of the adult bees in the colony about three times faster than that of fenitrothion though both pesticides provide the similar influence on capped brood. Therefore, we think that the extinction of the dinotefran colony was attributed to the rapid unbalancing of the number of worker bees in a colony.

We speculate the following negative influence of neonicotinoids on honeybee colonies in the natural environment from our field experimentals: Since a neonicotinoid is a tasteless, odorless and persistent pesticide, honeybees continue to take it for a long time from water in fields. For instance, a rice paddy is one of the typical water resources for bees in Japan. Since a persistent neonicotinoid is accumulated in the bodies of honeybees even if its concentration is much lower than that of our experiments, it influences in particular elder worker bees and causes a collapse of the colony maintained by the worker bees. On the other hand, the organophosphates are unstable and not persistent in toxicity, which may lead to a rapid decay of toxicity with time.

In this experiment, we did not confirm the perfect CCD caused by the both pesticides. Although an extinction occurred for dinotefran colony, many dead bees were found near the hive, which does not satisfy the condition of the CCD generation. The other aspects of CCD such as the existence of the queen, capped broods, and enough foods in the colony just before the extinction, however, were observed. We therefore think that a partial CCD occurred in the dinotefuran colony. In this field experiments, the water resource was placed with pesticides in their hives, which gives an artificial influence to the colony and may be a reason for observation of massive dead bees around their hive.

## Acknowledgments

This research was supported in part by Yamada Research Grant. Here, we would like to take this opportunity to thank all the individuals who assisted us in our research.

## Author Contributions

Toshiro YAMADA (TY), Yasuhiro YAMADA (YY) and Kazuko YAMADA (KY) conceived and designed the experiments. TY and KY performed the experiments. YY, TY and KY analyzed the data. Hiroko Nakamura (HN) counted the numbers of adult bees and capped brood on photos. TY and YY contributed reagents/materials/analysis tools. TY, YY and KY wrote the paper.

